# Welch-weighted Egger regression reduces false positives due to correlated pleiotropy in Mendelian randomization

**DOI:** 10.1101/2021.04.09.439229

**Authors:** Brielin C. Brown, David A. Knowles

## Abstract

Modern population-scale biobanks contain simultaneous measurements of many phenotypes, providing unprecedented opportunity to study the relationship between biomarkers and disease. However, inferring causal effects from observational data is notoriously challenging. Mendelian randomization (MR) has recently received increased attention as a class of methods for estimating causal effects using genetic associations. However, standard methods result in pervasive false positives when two traits share a heritable, unobserved common cause. This is the problem of correlated pleiotropy. Here, we introduce a flexible framework for simulating traits with a common genetic confounder that generalizes recently proposed models, as well as simple approach we call Welch-weighted Egger regression (WWER) for estimating causal effects. We show in comprehensive simulations that our method substantially reduces false positives due to correlated pleiotropy while being fast enough to apply to hundreds of phenotypes. We first apply our method to a subset of the UK Biobank consisting of blood traits and inflammatory disease, and then a broader set of 411 heritable phenotypes. We detect many effects with strong literature support, as well as numerous behavioral effects that appear to stem from physician advice given to people at high risk for disease. We conclude that WWER is a powerful tool for exploratory data analysis in ever-growing databases of genotypes and phenotypes.

## 1 Introduction

Modern population-scale biobanks contain genetic information with simultaneous measurements of many phenotypes, providing unprecedented opportunity to study the relationship between biomarkers and disease. However, inferring causal effects from observational data is notoriously challenging. Mendelian randomization (MR) has recently received increased attention as a class of methods that can mitigate issues in causal inference by using genetic variants (single nucleotide polymorphisms, SNPs) from genome-wide association studies (GWAS) as instrumental variables to determine the causal effect of an exposure (*A*) on an outcome (*B*). To estimate causal effects, MR methods must make strong assumptions that limit their ability to be applied at scale. Perhaps the most problematic assumption is that the SNP only effects B through A, *i.e.* there is no horizontal pleiotropy. Recent methods such as Egger regression and the mode-based-estimator are able to relax this assumption, instead assuming there is no correlated horizontal pleiotropy or modal pleiotropy, respectively [1, 2]. Correlated horizontal pleiotropy arises when both *A* and *B* share a common heritable factor (*U* in Figure 1a), resulting in genetic correlation between the traits in the absence of a causal effect. This kind of pleiotropy is both challenging to handle and thought to be pervasive, with computationally-intensive mixture models recently showing success at estimating causal effects in this setting [3, 4]. Another approach, the latent causal variable (LCV) model, is able to detect causality under arbitrarily-structured pleiotropy [5]. However, the quantity that LCV calculates is not interpretable as the causal effect size of A on B.

**Figure 1:**
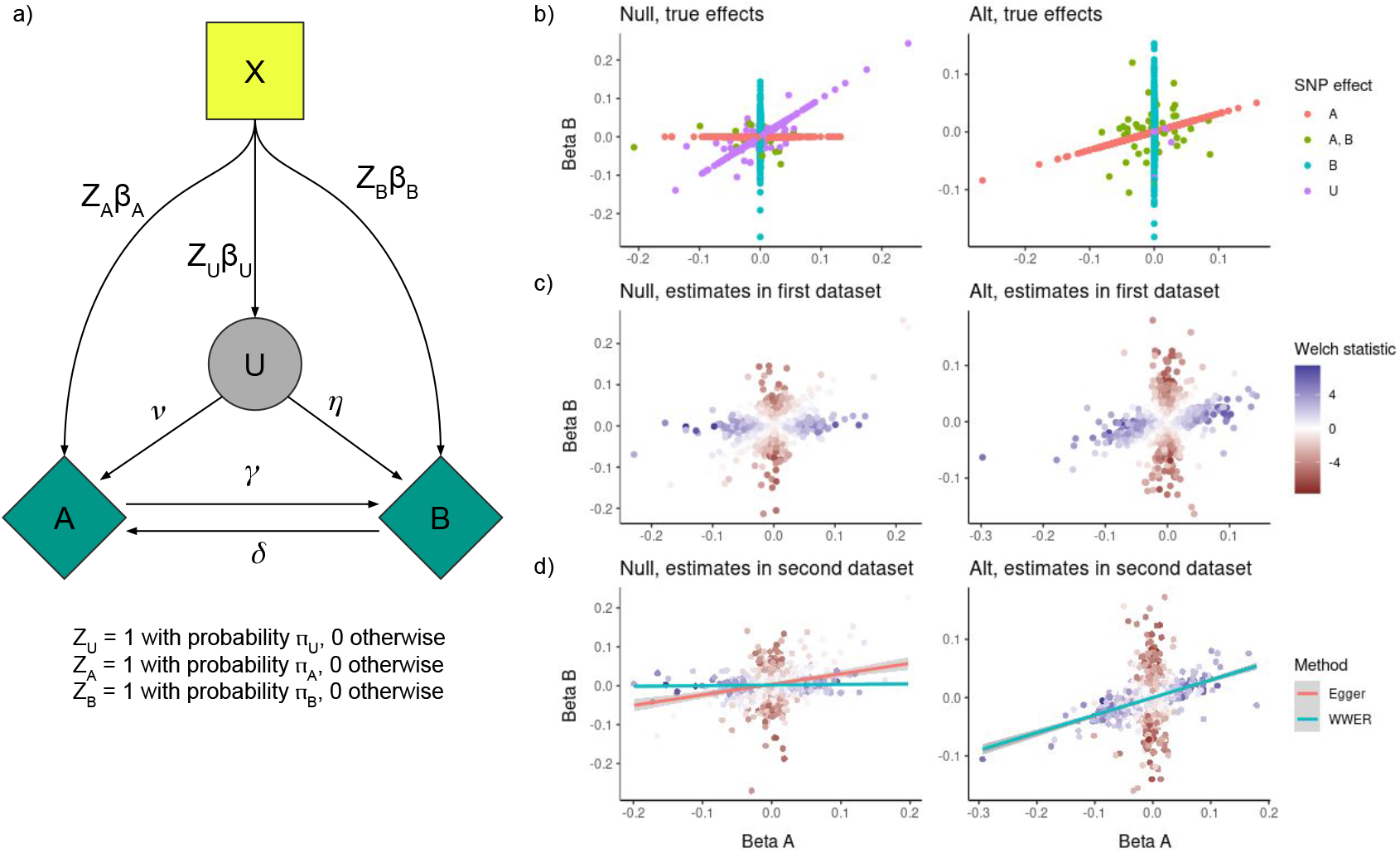
Our model for bi-directional Mendelian randomization, along with an example demonstrating the utility of WWER as compared to standard Egger regression under both the null and 1-way alternative hypothesis. a) A flexible model for bi-directional Mendelian randomization. SNPs *X* can effect the unobserved phenotype (confounder) *U* as well as the observed phenotypes of interest *A* and *B*. *η* and *ν* represent the per-variance effect of *U* on *A* and *B*, respectively, while *γ* and *δ* represent the per-variance causal effect of *A* on *B* and *B* on *A*, respectively. The allelic architecture of each phenotype can be independently adjusted by adjusting the proportion of effect variants, *π*, and variance of the distribution of effect sizes, *β*. b) The true effect of each SNP on phenotype A vs B under (left) a null model with 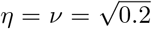 and *γ* = *δ* = 0 and (right) an alternative (alt) model with *η* = *ν* = *δ* = 0 and 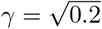. In the first sample, WWER calculates the Welch statistic, with large positive values (blue) indicating the SNP has a stronger effect on *A* and large negative values (red) indicating the SNP has a stronger effect on *B*. SNPs with near-equal effects on the diagonal axis get scores near 0. d) In the second sample, WWER filters SNPs with low Welch statistic, then uses the Welch statistic as a weight for the remaining SNPs when regressing the effect of the outcome on the exposure. Under the null (left) Egger regression produces a false positive, whereas WWER down-weights pleiotropic SNPs and does not. Under the alternative (right) both methods produce nearly-identical results.

Most MR studies also presuppose the direction of effect, specifying one phenotype as the outcome and the other as the exposure. Pre-specifying the effect direction can be sound when the outcome is clearly biologically downstream of the exposure, but many cases are less clear-cut and it is be preferable to learn the direction of the effect from the data. Some researchers have instead explored bidirectional MR [6, 7], which tests for an effect in each direction, or gwas-pw [8], which infers the effect direction from the data. Others have used Steiger filtering to remove instruments that might be acting on the outcome, rather than the exposure, which has been shown to reduce false positives due to misspecification of the exposure-outcome relationship [9]. However, the utility of these approaches for complex traits, which might contain non-causal correlated pleiotropy, is questionable [5].

Here, we introduce a flexible model for bi-directional MR that explicitly models the genetic architecture of both the observed phenotypes and a heritable confounder while allowing for arbitrary linear dependencies between them (Figure 1a). Our model captures recently proposed models for MR, including LCV [5] and CAUSE [3], as special cases. We also introduce a simple method for producing causal effect estimates that is based on filtering and down-weighting likely pleiotropic SNPs in an Egger-like regression, an approach we call Welch-weighted Egger regression (WWER, Figure 1b-d). By filtering SNPs with indistinguishable statistical effects on the exposure and the outcome, our method can be seen as an extension of Stieger filtering. To our knowledge Steiger filtering has not been extensively evaluated as an approach to dealing with non-causal association due to correlated pleiotropy. We show via extensive simulations varying the trait and model architectures that our approach reduces false positives due to correlated pleiotropy while being computationally efficient enough to apply in bi-directed exploratory data analyses of hundreds of phenotypes. We first apply our method to a limited set of phenotypes from the UK Biobank (UKBB) consisting of blood biomarker and blood cell composition traits, as well as common inflammatory diseases, and recover signals corresponding to known disease risk factors. We next apply our method broadly to over 400 phenotypes from the UKBB, again recovering known disease risk factors, while also finding broad signatures of risk factors on behavior, likely reflecting patient response to common medical advice.

## 2 Results

### Overview of methods

We introduce a flexible model that allows for both unidirectional and bidirectional causal effects while explicitly modeling the genetic architecture of each trait. In contrast to previously proposed models [5, 3], ours decouples the genetic architecture of the confounder from that of the exposure, allowing for arbitrary linear effects of the confounder on the pair of observed phenotypes. Our model is also agnostic to the labeling of either observed phenotype as the exposure or the outcome. In brief, we use *A* and *B* to denote the observed traits in the study, and *U* to denote the unobserved genetic confounder. SNPs *X* effect each of *A*, *B* and *U* with probabilities *q*, *r*, *s* and effect sizes *β_A_, β_B_, β_U_* sampled from a normal distribution with variances 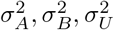, respectively. The probability of effect and variance of the sampling distribution combine to determine the genetic architecture of each trait independently of the others. Finally, *η* and *ν* specify the effect of the hidden confounder *U* on *A* and *B*, while *γ* and *δ* model the causal effect of *A* on *B* and *B* on *A*, respectively. Under this model, the phenotype values are given by

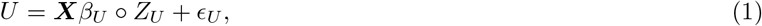

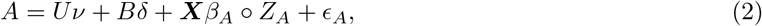

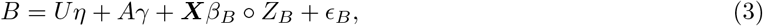

where *Z*’s represent indicator variables that the SNP effects that trait, sampled as indicated above, indicates vector element-wise (hadamard) multiplication, and bolding represents matrices. In Section 4 we show how to simulate from this model and parameterize it in terms of the heritability of each phenotype rather than the variance of the effect size distribution. We also explicitly describe how to set the parameters to mimic the models considered in [5] and [3]

To produce effect estimates, we introduce a simple method based on a modification to Egger regression [1] that down-weights likely pleiotropic SNPs. Similarly to Steiger filtering [9], we leverage the intuition that if *A* causes *B* and a SNP effects *A* directly, the per-variance effect of the SNP on *B* can be no larger than the per-variance effect of the SNP on *A* times the per-variance effect of *A* on *B*. That is, the SNP must have its per-variance contribution to *B* reduced by the effect of *A* on *B*. We use this to construct a novel weighting scheme for Egger regression. First, we select a *p*-value threshold *p_t_* (usually 5 × 10^*−*^8). For both phenotypes *A* and *B*, we construct a set of marginally associated SNPs at threshold *p_t_*. For this set of SNPs, we calculate a weight based on the Welch test statistic for a two-sample difference in mean with unequal variances, and the standard inverse-variance weight. If 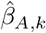 and 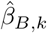 are our estimates of the effect of SNP k on phenotypes *A* and *B*, respectively, with 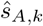 and 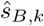 their standard errors, the Welch test statistic [10]

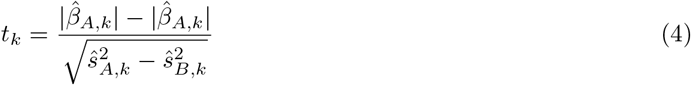

is and our weight is

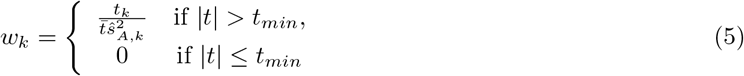

where 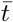 is the mean Welch statistic, and *t_min_* is the SNP inclusion threshold. We use these SNP weights in the Egger regression of *B* on *A* and vice versa for the reverse direction. To avoid bias, we must use two sets of summary statistics. The first set is for SNP selection and weight construction, and the second set is for estimating the causal effect. This method has two parameters, *p_t_* and *t_min_*. Here we choose not to tune them and instead always set them to *p_t_* = 5 × 10^−8^, corresponding to genome-wide significance, and *t_min_* = 1.96, corresponding to a two-sided *p*-value for a difference in mean effect of 0.05.

### Simulations

#### WWER reduces false discoveries due to correlated pleiotropy

Our first goal was to assess the calibration of WWER under the two-way null as compared to other methods under a broad range of simulation settings. In total we simulated 82 different combinations of simulation parameters. The first 20 settings are designed to mimic the simulations in [5] (Figure 2a), the next 20 settings are designed to mimic the simulations in [3] (Figure 2b), and the final 42 explore various combinations of high and low polygenicity for each of the observed and unobserved traits (Figure 2c). In all cases we evaluate the false positive rate (FPR) in both the *A* to *B* direction and the *B* to *A* direction. We also calculate the mean absolute error (MAE) of the effect size estimate. We compare WWER to the standard methods of inverse variance weighting (IVW) and Egger regression, as well as several more recently proposed methods: CAUSE [3], MR Mix [4], MR PRESSO [11], raps [12], the weighted median estimator (WME) [13] and the mode-based estimator (MBE) [2]. We also compare against Egger regression with Steiger filtering [9], which has not previously been evaluated for the purpose of handling correlated pleiotropy in bi-directional MR. We intentionally excluded methods such as gwas-pw [8] and LCV [5] that cannot produce bi-directed effect estimates.

**Figure 2:**
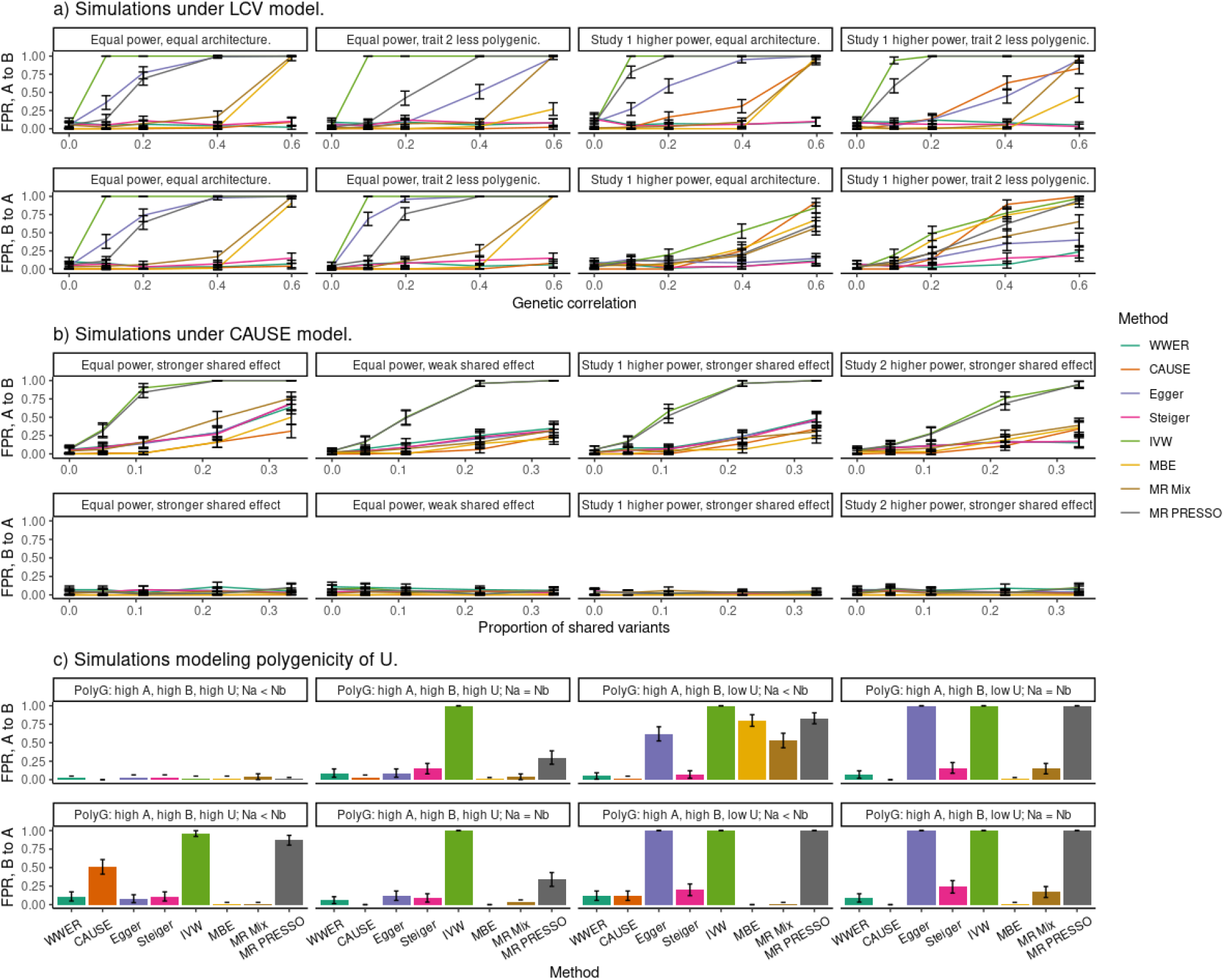
False positive rate in simulations under the bi-directional null for various settings of the simulation parameters. In all cases we consider both the *A* to *B* direction (top) and *B* to *A* direction (bottom). a) Simulations with parameters set to mimic the LCV model while varying the power and polygenicity of each trait (panels) as well as the genetic correlation (x-axis). WWER and Steiger filtering perform well, while other methods struggle with at least one setting. b) Simulations with parameters set to mimic the CAUSE model while varying the power and strength of the effect of the hidden node (*U*) on the observed traits *A* and *B* (panels). All methods with the exception of IVW and MR PRESSO perform well. c) Simulations explicitly modeling the polygenicity of *A*, *B*¡ and *U*, while varying the relative power of each study (panels). In the panels shown, there is a strong symmetric effect of the hidden node on the traits. Simulations with asymmetric effects are shown in Tables S8-9. There is no method that performs well in every setting, but WWER, Stieger filtering, CAUSE, the MBE, and MR Mix perform well overall. Our results are further summarized in Table 1.

The simulations corresponding to the LCV null model are broadly defined by 1) an equal effect of the hidden confounder (*U*) on both observed traits (*A* and *B*) and 2) a genetic architecture of *U* that results in an equal per-variance contribution of each SNP to *A* and *B* both when it acts directly on the them or through *U* (for more details, see Section 4). We evaluated settings where the studies for *A* and *B* had 1) equal sample sizes of 100,000 individuals and an equal genetic architecture, 2) equal sample sizes but trait *B* had half as many causal SNPs, 3) study *B* had reduced power (20, 000 individuals) and 4) study *B* had reduced power while being less polygenic. In every setting we simulate 500, 000 total SNPs, 2, 500 of which are causal for each observed phenotype. In each of these categories we varied *ν*, *η* and the proportion of shared causal SNPs such that the induced genetic correlation varied from 0.0 to 0.6. For complete settings for each simulation see Table S1. WWER and Egger regression with Stieger filtering maintained a low false positive rate across all of these simulation settings, while other methods had variable performance depending on the setting (Figure 2a). CAUSE also performed well unless the genetic correlation was 0.4 or above and the studies had unequal power, while MBE and MR-Mix also struggled with higher values of genetic correlation. Egger regression, MR PRESSO, IVW and raps all performed poorly overall. For space constraints we have omitted (r)aps and WME from Figure 2, for complete results see Tables S4-S5.

**Table 1:** A summary of the results from all of our null simulations. In each setting, we ranked every method according to its false positive rate (FPR) and mean absolute error (MAE). Then, we calculated the mean ranking of each method across all simulations settings (columns FPR rank and MAE rank, respectively). We also calculated the percentage of settings in which each method had *FPR <* 5%, indicating well-calibrated *p*-values, as well as the percentage of settings with *FPR <* 20%, indicating a controlled level of excess false positives (columns FPR < 5% and FPR < 20%, respectively. Finally, we calculated the time required to calculate an effect in each direction with each method (column Runtime).

| Method | FPR rank | MAE rank | FPR < 5% | FPR < 20% | Runtime (s) |
| --- | --- | --- | --- | --- | --- |
| WWER | 6.034 | 6.735 | 77.4 | 92.7 | 0.634 |
| Steiger | 5.329 | 7.372 | 76.2 | 92.1 | 0.634 |
| CAUSE | 4.210 | 4.671 | 81.7 | 90.9 | 2958.892 |
| MBE | 3.351 | 5.823 | 84.8 | 89.6 | 38.140 |
| MMR Mix | 4.125 | 6.134 | 76.8 | 87.2 | 79.670 |
| Egger | 6.683 | 9.183 | 59.8 | 67.1 | 0.632 |
| WME | 6.372 | 5.177 | 52.4 | 63.4 | 8.594 |
| MR PRESSO | 8.287 | 4.366 | 45.7 | 54.3 | 313.527 |
| raps | 8.381 | 6.421 | 39.0 | 50.0 | 0.684 |
| IVW | 9.418 | 6.683 | 36.0 | 40.2 | 0.640 |
| aps | 9.290 | 8.116 | 35.4 | 39.6 | 0.635 |

The simulations corresponding to the CAUSE model are broadly defined by 1) a much stronger effect of *U* on the exposure (*A*) relative to the outcome (*B*) and 2) a genetic architecture of the hidden trait that results in an equal per-variance contribution to the *A* whether the SNP acts directly on it or via *U* (again see Section 4 for more details). Since our model does not make an exposure/outcome distinction, we choose to use *A* as the exposure and *B* as the outcome, but we continue to evaluate performance in both directions. In all settings we simulated 500, 000 SNPs with 1, 000 causal SNPs per observed phenotype. We chose four broad categories wherein we adjusted the proportion of the causal variants effecting *U* from 0.0 to 0.33. The four categories correspond to 1) equal power (100, 000 individuals) with a stronger shared effect (explaining 5% of the variance in *B*), 2) equal power with a weaker shared effect (explaining 2% of the variance in *B*), 3) study 2 having lower power (20, 000 individuals) with the stronger shared effect and 4) study 1 having lower power (20, 000 individuals) with the stronger shared effect. For complete settings for each simulation see Table S2. In this setting, CAUSE, all Egger-based methods as well as MR Mix and the MBE perform similarly well with a well-controlled error rate at lower proportions of shared variants and some excess false positives at higher levels. The WME, MR PRESSO, and (r)aps perform similarly to IVW which struggles to control false positives even for relatively small fraction of pleiotropic SNPs in all settings. CAUSE seems to perform better here than in similar situations in the original manuscript [3]. This is likely because we are using pre-pruned variants without linkage disequilibrium (LD), unlike in [3] where CAUSE must additionally handle LD. In the *B* to *A* direction, all methods are able to control the false positive rate. For complete results see Tables S6-S7.

Finally, we exhaustively tested all combinations of 1) low (500 directly causal variants) and high (2000 directly causal variants) polygenicity for each of *A*, *B*, and *U*, 2) either equal (100,000 individuals per study) or unequal (25,000 individuals in the under-powered study) sample sizes, and 3) either equal effects of *U* on both traits (explaining 30% of the variance in each) or unequal effects (explaining 30% in one and 10% in the other). For complete simulation settings see Table S3. Due to space limitations we present 8 of the 42 resulting combinations in Figure 2 and the rest in Tables S8-S9. Perhaps unsurprisingly, some settings favor certain methods over others and there is no method that controls the FPR in every setting. WWER and Egger regression with Steiger filtering worked well for most settings. However, there were notable settings where these methods performed poorly. For example, when the polygenicity of *A* is high, the polygenicity of *U* is low, *U* has a larger effect on *A*, and the sample size of *A* is large, both methods produce a false positive > 95% of the time. (Table S8 lines 5 and 33, Table S9 lines 2 and 16). Another interesting setting is low polygenicity of both *A* and *B*, high polygenicity of *U*, equal sample sizes and a larger effect of *U* on the exposure (Table S9 line 14). In this setting WWER produces many excess false positives (*FPR* = 0.47 ± 0.05), but this is about half as many as standard Egger regression or Egger with Steiger filtering (*FPR* = 0.88(0.03) and *FPR* = 0.80(0.04), respectively). With some exceptions, CAUSE and MBE also performed well in these settings, while all other methods performed poorly overall.

We summarize our results in Table 1 by ranking the methods according to FPR and mean absolute error (MAE) in each of the simulations settings in both directions. We also calculate the percentage of settings where each method had an estimated *FPR* whose 95% confidence interval contained either 0.05, indicating well-calibrated *p*-values, or 0.20, indicating that the excess false positives are limited to a useful level. The MBE, CAUSE, and MR-Mix performed quite well overall. These three methods generally produced the best-calibrated *p*-values and controlled the *FPR* at the 0.05 level in the highest percentage of tested cases. However, WWER followed by Egger with Stieger filtering produced a controlled amount of excess false positives (*FPR* < 20%) more frequently than other methods. WWER generally produced slightly less conservative *p*-values while having a lower MAE overall as compared to Egger with Stieger filtering. While these two methods did perform similarly, as mentioned above we found evidence of cases where WWER out-performed Egger with Stieger filtering by a large margin, but the opposite was never true (Tables S8-S9). A final consideration, especially in exploratory data analysis applications, is run-time. Regression-based methods are very fast, while more sophisticated methods can take much longer. CAUSE took nearly 50 minutes on average to calculate effects in both directions (Table 1). While the MBE and MR-Mix are somewhat faster, we used only a small number of sampling iterations (1000) to generate *p*-values accurate to two significant digits. In exploratory data analysis cases, where the multiple testing burden is likely to be high, many more iterations will need to be used to generate *p*-values with more significant digits.

#### WWER maintains power

Our next goal was to evaluate the power of WWER as compared to other methods when the alternative hypothesis is true and there is no uncorrelated pleiotropy (*γ* and/or *delta* ≠ 0, while *ν* = *η* = 0). We considered both the unidirectional (*A → B*, *γ* > 0, *δ* = 0) and the bi-directional (*A ⟷ B*, *γ* ≠ 0, *δ* = 0) alternate while varying the strength of the effects (*γ* or *δ*), the polygenicity of *A* and *B*, and the sample size of each study (Figure 3).

**Figure 3:**
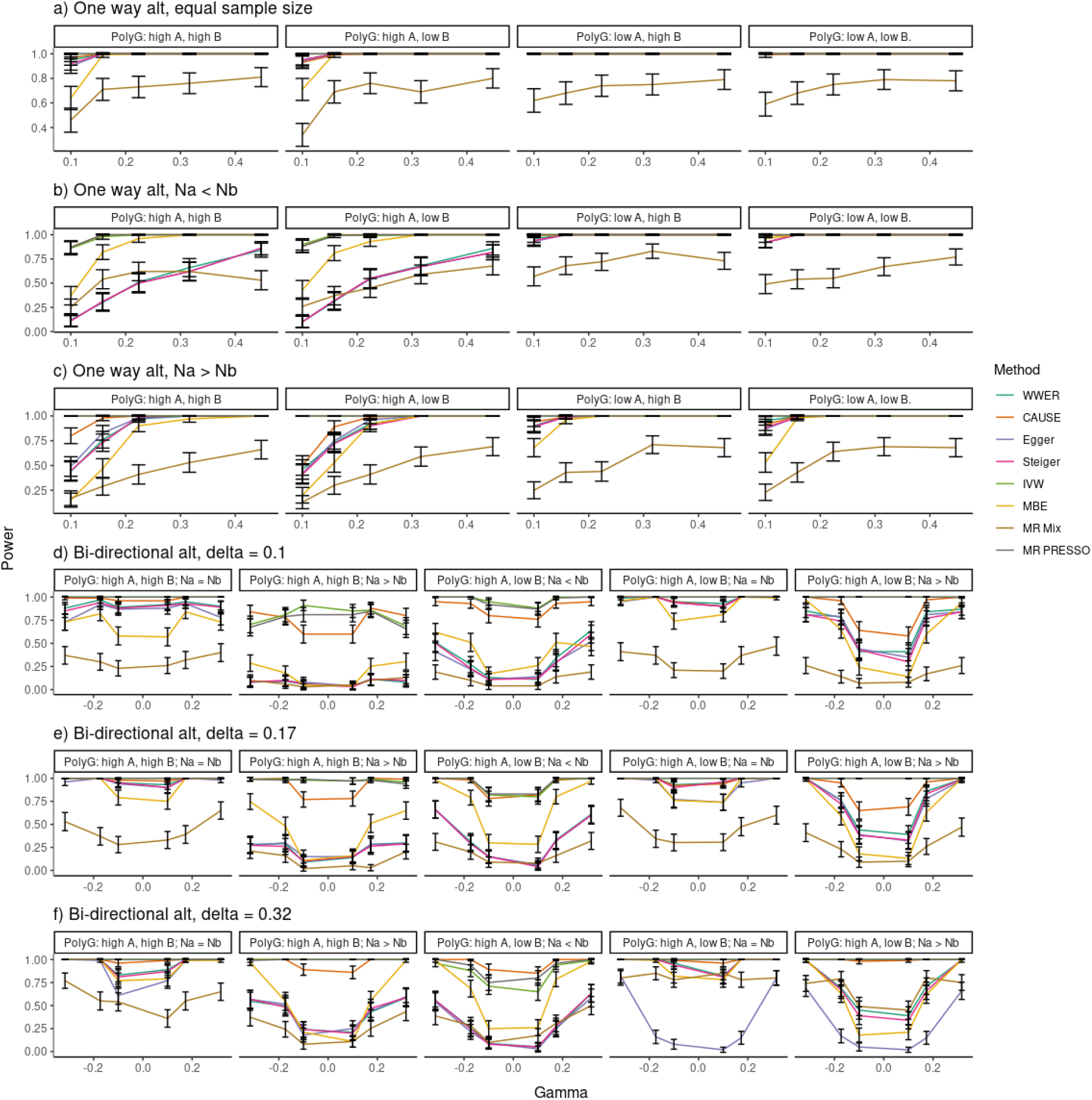
Power analysis of simulations under both the one way (A causes B, a-c) and bidirectional (A causes B and B causes A, d-f) alternative, without additional pleiotropy. a) With equal sample sizes, all methods except MR Mix show high power for all settings of the polygenicity of A and B (panels). b) When study A has lower power and the polygenicity of A is higher, regression-based methods have reduced power and are out-performed by the MBE. c) When study A has higher power, the opposite is true. (d-f) The power to detect an effect in both directions for all combinations of polygenicity and power as a function of the effect of A on B for various values of the effect of B on A.

In our simulations under the unidirectional alternative (Figure 3a-c), we varied the proportion of variance in *B* explained by *A* from 1% 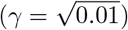 to 20% 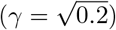. In our first simulation, both studies had equal power and we additionally tested high (2000 directly causal variants) and low (500 directly causal variants) polygenicity with a heritability of 25%. For complete simulation settings see Table S10. WWER and the other regression methods Egger regression and Stieger filtering performed similarly well, with generally strong performance for effect sizes above 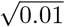 but reduced performance when *A* was highly polygenic but its study was under-powered. MR PRESSO and CAUSE show generally improved power in these more difficult cases, while the MBE can improve power when the study for *A* is under-powered but reduce it when the study for *B* is under-powered. MR Mix generally performs poorly compared to the other methods. Complete results are given in Table S13.

In our simulations under the bidirectional alternative (Figure 3d-f),we evaluated power to detect both effects (*A* → *B* and *B* → *A*) simultaneously. We set *δ* to either 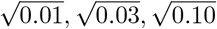 and varied *γ* from 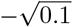 to 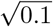 and again adjusted the polygenicity of each trait and the sample size of each study. For complete simulation settings see Table S12. Many trends from the unidirectional alternative were replicated here. Specifically, CAUSE, IVW and MR PRESSO performed well overall. The regression based methods performed similarly well in most settings, with lower power when the sample-sizes were unequal. The MBE was generally out-performed by regression based methods when the studies had equal sample size, but the opposite was true for unequal sample sizes, especially when the effects were larger. MR Mix again had poor power overall. Interestingly, we found two settings where standard Egger regression was substantially out-performed by both WWER and Egger with Steiger filtering: 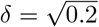, high polygenicity of *A*, low polygenicity of *B*, and either equal sample sizes or a larger sample size for *A* (Figure 3d-f). Complete results are given in Table S17.

Since we are concerned with estimating effects in both directions, we must take care to verify that under the unidirectional alternative, high power in the *A* to *B* direction (alternative hypothesis is true) does not result in a high false positive rate in the *B* to *A* direction (null hypothesis is true). In Figure S1 we plot the *FPR* in the *B* to *A* direction as a function of *γ* for each corresponding simulation in Figure 3a-c. All methods except IVW, standard Egger regression, and MR PRESSO were able to control the reverse false positive rate. This was primarily an issue for larger effects 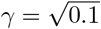 and 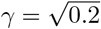 when study *B* had high power. Complete results are given in Table S14.

Finally, we considered the one-way alternative hypothesis in the presence of correlated pleitropy (*γ, η, ν* > 0). In this setup both studies had equal power and we varied the strength and symmetry of the effect of *U*, considering equal weak pleiotropy 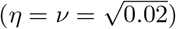, equal stronger pleiotropy 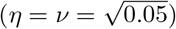, or unequal pleiotropy with a stronger effect on either *A* or *B*. For complete simulation parameters see Table S11. On the one hand, pleiotropy increases the power to detect the non-null effect because it lends additional signal supporting the effect of *A* on *B* (Figure S2). On the other hand, it also leads to additional false positives in the reverse direction (Figure S3). Here MR PRESSO and IVW produce a false positive nearly all the time if there is a strong pleiotropic effect on *A*. Standard Egger regression performs worse here than in the simpler setting of a true alternative hypothesis with no pleiotropy, but WWER and Stieger filtering are able to reduce this false positive rate substantially. Here, WWER clearly out-performs Steiger filtering in settings where they both produce excess false positives, such as when the polygenicity of *A* and *B* are high but the polygenicity of the confounder is low. For complete results see Table S15-16.

#### Application to blood traits and immune disorders in the UK Biobank

There are a number of common disorders involving immune system and inflammatory response disregulation (immune mediated inflammatory disease, IMID), such as allergy, asthma, diabetes and psoriasis, among others [14]. Blood is both an easily accessible tissue and a heterogeneous mixture of numerous cell types with relevance to inflammatory and immune response, so there is a strong interest in intermediate blood biomarkers of IMIDs for measuring disease risk, monitoring progression, and developing treatments [14, 15]. The UK Biobank contains measurements of clinical laboratory biomarkers, as well as blood cell-type composition and disease phenotype data for > 480, 000 individuals [16]. We obtained summary statistics for sex-split UK Biobank phenotypes from the Neale lab, who corrected for age, age^2^ and 20 principal components of the genotype matrix [17]. For ease of interpretation, we transformed all effect sizes to the per-variance scale. We filtered for phenotypes with an LD score regression heritability Z-score above 4 and at least “medium” confidence, and removed one phenotype from every pair with genetic correlation above 0.9 to avoid including what are effectively duplicate traits. We used male summary statistics for SNP selection and weight estimation, and female summary statistics for effect estimation. We removed any trait with an estimated male-female genetic correlation *ρ_g_ <* 0.5 or a Z-score for non-zero genetic correlation below 2. This left us with 21 measurements of blood composition, 20 blood biomarkers and 10 IMIDs (Table S18). We used LD-pruned SNPs attaining genome-wide significance (*p* ≤ 5 × 10^−8^) as instruments with WWER to estimate causal effects (CE) of each biomarker on each disease, and vice-versa (disease on biomarker).

We found 83 (of 410) significant effects at FDR 5% in the biomarker to disease direction after accounting for multiple testing using the Benjamini-Hochberg (BH, we denote adjusted *p*-values with *q* in the following) procedure (Figure 4a, Table S19). We observed a strong effect of platelet traits on asthma and allergy. For example, increased platelet distribution width (PDW) decreases asthma risk (*CE* = −0.034*, q* = 4 × 10^−10^) and allergy risk (*CE* = −0.016*, q* 2 × 10^−2^), increased mean platelet volume (MPV) decreases asthma risk(*CE* = −0.014*, q* = 2 × 10^−2^) and increased platelet*-*crit decreases allergy risk (*CE* = −0.066*, q* = 3 × 10^−10^). Platelet traits have long been implicated in asthma and allergy [18, 19, 20], with lower MPV values observed in individuals with asthma and allergy and lower PDW values observed in individuals with asthma. Platelet traits are now thought to play an important role in both the innate and adaptive immune response [21]. We find that PDW is implicated in 7 of the 10 IMID studied, and MPV is implicated in 4 of the 10. This gives evidence that platelet activity can have an effect on immune-system function, with broad downstream consequences that include many common diseases.

**Figure 4:**
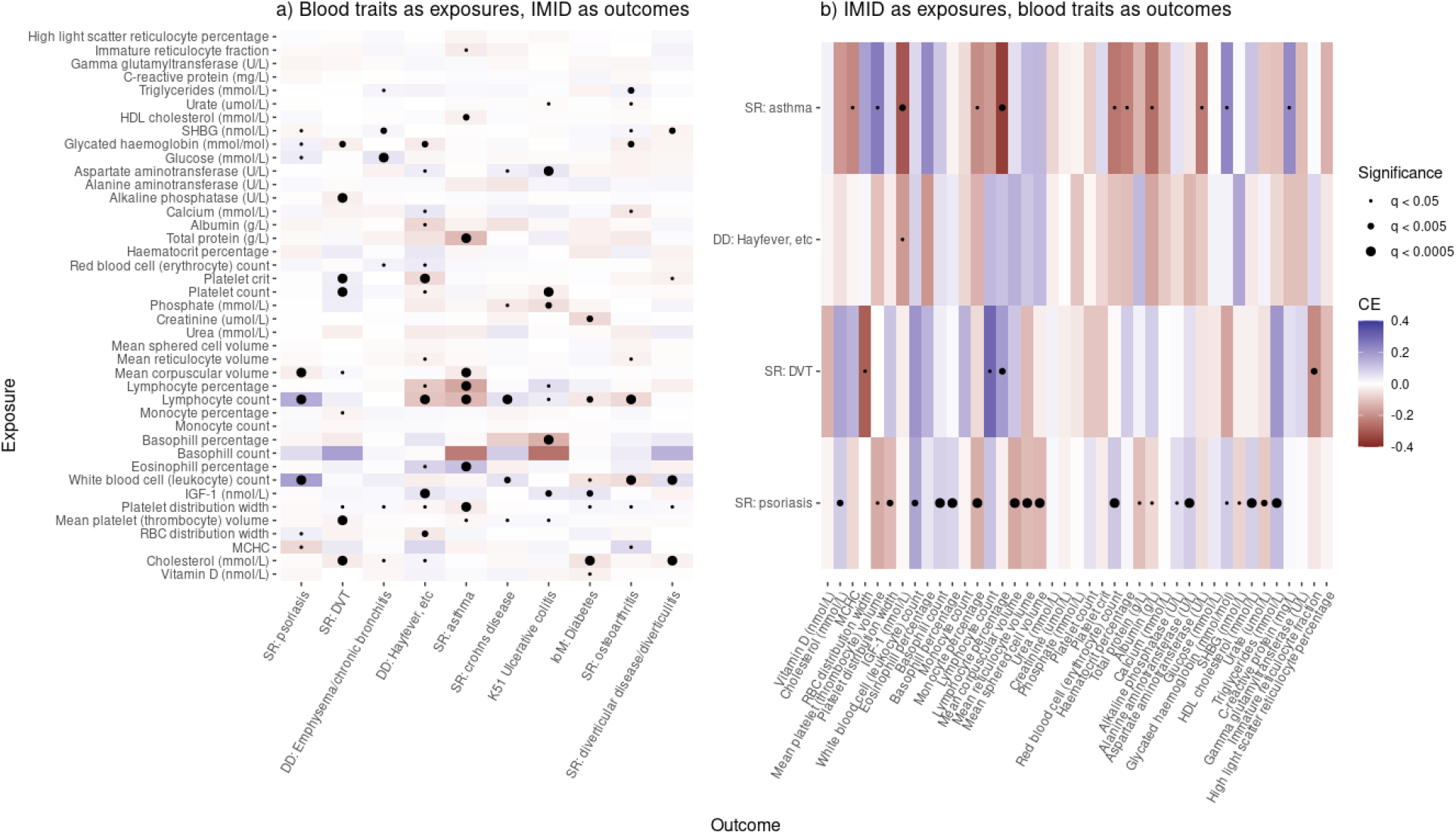
An investigation into the relationship between immune-mediated inflammatory diseases and blood biomarkers in the UK Biobank. (a) Estimated causal effects using blood traits as exposures and IMID as outcomes replicates known disease biology. (b) Estimated causal effect using IMID as exposures and blood traits as outcomes reveals many significant “reverse” causal effects. Dots indicate level of statistical significance of *p <* 0.05 after FDR correction.

Lymphocyte count, a marker of inflammation, is also implicated in 7 of the 10 IMID that we analyze. We detect effects of increased lymphocyte count on psoriasis (*CE* = 0.159*, q* = 1 × 10^−9^), Crohn’s disease (*CE* = 0.057*, q* = 3 × 10^−5^), and ulcerative colitis (*CE* = 0.037*, q* = 4 × 10^−2^). We detect effects of decreased lymphocyte count on asthma (*CE* = −0.12*, q* = 6 × 10^−9^), osteoarthritis (*CE* = −0.069*, q* = 4 × 10^−6^), allergy (*CE* = −0.101*, q* = 3 × 10^−5^) and diabetes (*CE* = −0.04*, q* = 3 × 10^−3^). A lower neutrophil to lymphocyte ratio has been observed in patients with each of these diseases [22, 23, 24, 25]. In several of these results, our estimated CE and the genome-wide genetic correlation have opposite signs. For example, the genetic correlation between lymphocyte count and asthma is positive (*ρ_g_* = 0.054), as is the genetic correlation between lymphocyte count and osteoarthritis (*ρ_g_* = 0.212). In each of these cases, the negative effect direction inferred by WWER is more consistent with the observed lower neutrophil to lymphocyte ratio in these diseases. This indicates that the total genetic correlation can be misleading, even in the presence of a causal effect, it is possible for a genetic confounder, or possibly random noise, to result in an observed genetic correlation with a different sign than the true causal effect.

Total cholesterol also has several disease consequences. We observe protective effects of increased total cholesterol level on diabetes (*CE* = −0.047*, q* = 4 × 10^−10^), deep vein thrombosis (DVT, *CE* = −0.035*, q* = 3 × 10^−8^), diverticulitis (*CE* = −0.025*, q* = 5 × 10^−4^), and emphysema (*CE* = −0.016*, q* = 4 × 10^−2^). We also observe a protective effect of increased HDL cholesterol level on asthma (*CE* = −0.026*, q* = 2 × 10^−3^) These findings are particularly interesting in light of recent work suggesting that cholesterol can lower inflammation [26], that higher cholesterol is a consequence of the body’s attempt to control inflammation, rather than the cause of disease in itself [27]. Interestingly we observe a weak effect of increased cholesterol on allergy risk(*CE* = 0.021*, q* = 4 × 10^−2^) which is inconsistent with the genetic correlation between these traits (*ρ_g_* = −0.014). Cholesterol is known to effect development of allergy, but reports differ on the direction of the effect [28].

Other notable effects we observe include a strong effect of eosinophil percentage on asthma (*CE* = 0.118*, q* = 5 × 10^−6^), aspartate aminotransferase on ulcerative colitis (*CE* = 0.047*, q* = 5 × 10^−5^), glucose on emphysema (*CE* = 0.051*, q* = 9 × 10^−5^), and a protective effect of vitamin D on diabetes (*CE* = −0.024*, q* = 5 × 10^−2^). Eosinophils are known to play an important role in the pathogenesis of asthma [29], with well-established genetic evidence indicating a protective effect of lower eosinophil count on asthma risk [30]. Liver test abnormalities are frequently observed in patients with inflammatory bowl diseases [31] and appear to be an risk factor for complications in patients with Crohn’s disease [32]. Blood glucose has been observed to be elevated in patients experiencing chronic obstructive pulmonary disease (COPD) exacerbations [33]. Vitamin D has been linked to the onset of diabetes [34].

We found 36 (of 164) significant effects in the disease to biomarker direction after accounting for multiple testing using the BH procedure (Figure 4b, Table S20). Most of these are driven by just two phenotypes: 20 are effects of psoriasis and 11 are effects of asthma. Some of the top effects of psoriasis are related to red blood cells (RBC). We estimate that psoriasis decreases mean sphered cell volume (*CE* = −0.12*, q* = 1 × 10^−7^), mean corpuscular volume (*CE* = −0.15*, q* = 1 × 10^−7^), and mean reticulocyte volume (*CE* = −0.12*, q* = 4 × 10^−5^), while increasing red blood cell count (*CE* = 0.12*, q* = 1 × 10^−7^). There is an established relationship between red blood cell function and psoriasis [35, 36, 37]. There is disagreement in the literature about the correlation of red blood cell count and psoriasis, with one study showing an increase in patients, consistent with our results [37] and others showing a decrease, inconsistent with our results [35, 36]. However, the latter study also shows that treatment of psoriasis can correct RBC damage, which might suggest that psoriasis is the cause, rather than consequence, of RBC damage. We also observe effects of psoriasis on lipid profile. We infer psoriasis increases HDL cholesterol (*CE* = 0.088*, q* = 3 × 10^−4^), total cholesterol (*CE* = 0.097*, q* = 2 × 10^−3^), and triglycerides (*CE* = 0.123*, q* = 4 × 10^−7^). Psoriasis is well-known to be co-morbid with cardiovascular disease, and dislipidemia has long been observed in patients with psoriasis [38, 39]. However, we note that our inferred direction of effect for HDL cholesterol is inconsistent with prior literature showing decreases in HDL cholesterol level in patients with psoriasis, and with the genetic correlation between the traits (*ρ_g_* = − 0.21). Psoriasis is known to have a complex effect on HDL cholesterol function [40], and it is likely that the genetic instruments we use to estimate this effect on serum HDL levels do not reflect the complexity of this interaction.

Inferred effects of asthma include decreases in IGF-1 (*CE* = −0.308*, q* = 1 × 10^−3^), lymphocyte percentage (*CE* = −0.354*, q* = 2 × 10^−3^), and monocyte percentage (*CE* = −0.205*, q* = 5 × 10^−2^), and increases in C-reactive protein (*CE* = 0.226*, q* = 2 × 10^−2^) and glycated haemoglobin (*CE* = 0.233*, q* = 2 × 10^−2^). Glycated haemoglobin and C-reactive protein have both been observed to be elevated in patients with asthma [41, 42]. Monocytes and lymphocytes are both known to play an important role in asthma [43, 44, 45], however it is unclear how the impact of recruitment of specific monoctye and lymphocyte subsets to the lungs in asthma patients would impact circulating blood levels of these broad cell types. IGF-1 is known to play a function in the repair of lung tissue [46]. Serum IGF-1 level is known to be anti-correlated with asthma incidence and severity in the UK Biobank [47]. Our results suggest this is a consequence rather than a cause of asthma.

#### Phenome-wide analysis in the UK Biobank

The simplicity and speed of our method allows it to easily scale to phenome-wide analysis. Therefore, we obtained summary statistics for the remaining sex-split UK biobank phenotypes from the Neale lab [17], and applied the same pre-processing and phenotype selection filters used in the previous section. This left us with 411 phenotypes (Table S21). Of the 411 phenotypes chosen for analysis, 153 had at least 5 independent GWAS significant loci (Table S21). We used WWER to estimate the CE of all 153 phenotypes with at least 5 GWAS-significant loci on all 411 phenotypes. This results in bi-directed effect estimates for the 11, 628 pairs of traits where both have at least 5 instruments, and uni-directional effect estimates for the remaining 39, 474 pairs for a total of in 62, 730 CE estimates. Of these, we found 5, 770 effects (9.2%) were significant at a 5% FDR. Complete results for all tested pairs of phenotypes are given in Table S22.

We were curious to compare our CE estimates against estimates of genetic correlation in the same dataset. First, we clustered phenotypes by genetic correlation to determine if the patterns observed are shared in the CE estimates. While there are some similar patterns across the two matrices, the structure in the CE estimates is not as well-defined (Figure 5). Indeed, we find that while the CE estimates and genetic correlation estimates are correlated, that correlation is fairly weak (*r* = 0.175± 0.004). This weak correlation seems to be driven by CE estimates with large standard error. Accordingly, restricting our analysis to CE estimates with standard error below 0.05 yields a much stronger correlation (*r* = 0.573 ± 0.005). In general we found that more significant CE estimates were more similar to estimates of genetic correlation (Figure S4). As expected, the presence of genetic correlation does not indicate a detectable CE, and the causal effect and the total genetic correlation need not even have the same sign. However, strong CEs do frequently result in a total genetic correlation of similar magnitude.

**Figure 5:**
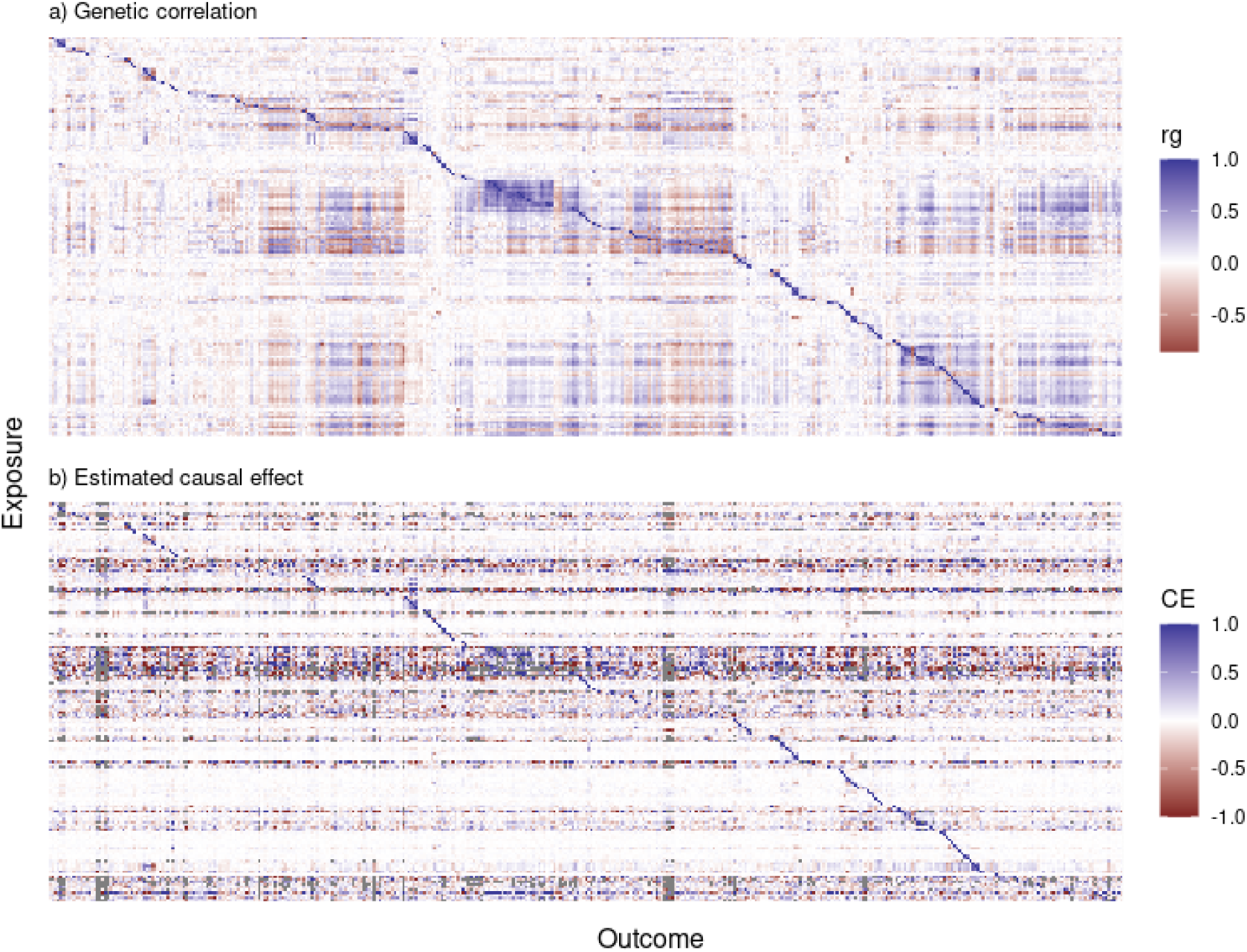
A comparison of genetic correlation (a) with the estimated causal effect (b). We calculated causal effects for all pairs of phenotypes passing our inclusion criteria using LD-pruned GWAS-significant variants as instruments. If both traits did not have significant variants, we calculate a unidirectional effect where the trait with significant variants is the exposure, whereas if both traits have significant variants we calculate a bidirectional estimate. Gray entries in (b) indicate pairs where the exposure had no remaining instruments after filtering likely pleiotropic SNPs, resulting in an NA value. Phenotypes are arranged by clustering on genetic correlation of traits that remain as exposures.

There were several traits with numerous consequences. The top 5 were white blood cell count (WBCC) with 188 effects, cholesterol with 173 effects, lymphocyte count with 172 effects, sex-hormone binding globulin with 154 effects, and body mass index with 147 effects. The top consequence of higher WBCC was an increase in “nervous feelings” (*CE* = 0.12*, q* = 1 × 10^−16^). WBCC is known to be elevated in individuals with depression and anxiety [48], and could reflect an effect of systemic inflammation on mood. The next strongest effect was a decrease in whole body water mass (*CE* = −0.236*, q* = 1 × 10^−16^). While dehydration is well-known to cause elevated WBCC, our results suggest that the opposite may also be true - higher levels of circulating WBCC could cause the body to retain less water. Two other strong effects of WBCC are on morphology, with an increase in WBCC resulting in a decrease in hip circumference (*CE* = −0.179*, q* = 2 × 10^−16^) and sitting height (*CE* = −0.23*, q* = 9 × 10^−16^). One study of Japanese men found that height and WBCC were inversely correlated, and concluded that this association may result from the presence of inflammation [49]. Interestingly, several of the top consequences of high cholesterol seemed to reflect behavioral changes resulting from common medical advice. For example, we found an effect on increased cholesterol on decreased use of butter (*CE* = −0.096*, q* = 1 × 10^−16^) and increased use of “other spread/margerine” (*CE* = −0.09*, q* = 1 × 10^−16^). We also found increased cholesterol caused a decrease in “salt added to food” (*CE* = −0.048*, q* = 1 × 10^−16^), and a decrease in “major dietary changes in the last year: no” (*CE* = −0.069*, q* = 1 × 10^−16^), indicating high cholesterol results in broad dietary changes. This phenomenon extends to choice of pain medication. We detect a positive effect of high cholesterol on aspirin use (*CE* = 0.048*, q* = 8 × 10^−13^) and a negative effect on ibuprofen use (*CE* = −0.026*, q* = 5 × 10^−5^). This is likely to reflect common medical advice for patients at risk of heart disease to choose aspirin, which has long been thought to reduce risk [50], and avoid ibuprofen, which is thought to reduce the effectiveness of aspirin [51]. We also replicate cholesterol as a known risk factor for heart disease (*CE* = 0.086*, q* = 1 × 10^−16^), which likely also accounts for an observed effect of high cholesterol on earlier “fathers age at death” (*CE* = −0.069*, q* = 1 × 10^−16^).

Several of the top consequences of body mass index (BMI) were also behavioral. For example, we observed a negative effect of BMI on using semi-skim milk (*CE* = −0.24*, p <* 1 × 10^−16^), but a positive effect on using skim milk (*CE* = −0.305*, p <* 1 × 10^−16^). We also observe a negative effect of higher BMI on “major dietary changes in the last year: no” (*CE* = −0.236*, p <* 1 × 10^−16^) and a positive effect on “major dietary changes in the last year: yes, because of other reasons [than illness]” (*CE* = 0.206*, p <* 1 × 10^−16^). These could again reflect behavioral consequences of common medical advice. Other effects of BMI were on blood biomarkers. For example, we observed an effect of higher BMI on higher C reactive protein (CRP, *CE* = 0.353*, p <* 1 × 10^−16^), lower albumin (*CE* = −0.222*, p <* 1 × 10^−16^), and higher urate (*CE* = 0.383*, p <* 1 × 10^−16^). Higher BMI is well known to cause higher serum urate levels [52], adipose tissue is known to induce low-grade inflammation, which can be measured by elevated CRP levels [53], and BMI is a known risk factor for hypoalbuminemia [54]. We find the known effect of BMI on diabetes (*CE* = 0.195*, p <* 1 × 10^−16^), but also find that BMI has broad effects on health and results in a lower “overall health rating” (*CE* = 0.149*, p* = 7 × 10^−8^).

## 3 Discussion

We have introduced a model for bi-directional Mendelian randomization with correlated pleiotropy that allows for flexibility in the specification of the genetic architecture for each trait, as well as a simple method for estimating causal effects called Welch-weighted Egger regression (WWER). We have shown that our method reduces false positives due to correlated pleiotropy compared to traditional methods in a broad range of simulation settings that encompass other recently-proposed models, and is fast enough to be applied at-scale.

We first applied WWER to a subset of the UK biobank comprising blood biomarkers and inflammatory disorders, and then more broadly to all heritable phenotypes in the biobank. Our initial analysis reiterated the role of platelet traits in the pathogenesis of asthma and allergy, and found that cholesterol and white blood cell count contribute broadly to inflammatory disease, among other findings. Our broad analysis found thousands of causal effects, many of which stem from a handful of broadly-impactful phenotypes. We replicate several known risk factors for disease such as high cholesterol on heart disease and high BMI on diabetes, but also detect numerous behavioral changes that seem to result from common physician advice.

Our approach builds on recent MR literature. By filtering genetic variants that have a statistically indistinguishable effect on both the exposure and the outcome, our method is closely related to Steiger filtering [9], which was conceptualized as a method for inferring the effect direction and has not received attention as a method for controlling for correlated pleiotropy. The primary conceptual difference is that we use the test statistic as a regression weight when calculating the effect of the exposure on the outcome with the retained SNPs. Compared to Steiger filtering, we control the FDR in slightly more of the tested settings, while also producing estimates with a lower mean absolute error. There are a small number of settings in which WWER produces a much lower false positive rate than Stieger filtering, but the reverse is never true. However, we find that both methods are generally useful for controlling for correlated pleiotropy. Our approach can be viewed a simple heuristic for classifying variants as effecting the exposure, the outcome, or the hidden variable. More sophisticated mixture model based methods, such as MR-mix and CAUSE, are also based on fitting the causal effect using a subset of SNPs that appear to effect the exposure. While these methods also work well in our simulations, they can take a prohibitively long time to run, preventing their application at the scale considered here. By removing genetic instruments with ambiguous effects, our method sometimes filters all potential instruments and cannot estimate the effect. We view this as both an advantage and a disadvantage: we avoid estimating an effect in ambiguous cases, but cannot always produce an estimate.

Despite its advantages, our approach has several limitations. First, our method requires that we split the initial cohort into instrument discovery and effect estimation sub-cohorts. While this approach is common in MR methods that must first identify instruments, this reduces power and two sets of summary statistics are not always available. Other recent approaches, such as CAUSE and LCV, have the advantage of modeling the entire spectrum of SNP-trait associations. Second, while our method reduces excess false positives, it does not completely eliminate them. Therefore, a small but notable number of statistically significant results in any large-scale analysis may be due to correlated pleiotropy. We have shown that these failure cases usually correspond to situations where the hidden factor has a strong effect on the exposure, and the exposure does not have many independent large-effect instruments. In this setting, the genetic signature of the exposure and hidden variable are difficult to distinguish. However, the fact that the hidden trait is highly causal for the exposure indicates that these cases may still be biologically interesting, even if they are not directly causal. One advantage of our method is that it only requires GWAS summary statistics, which are both legally and practically easier to share, and faster to work with when the primary data is large [55]. However, summary statistics are inherently limiting. Their use relies on the assumption that the creator of the summary statistics properly controlled for the relevant factors, which may not always be the case when the data are curated by groups without specific expertise in each of the relevant phenotypes. A final limitation of our method is that it estimates the total effect of the exposure on the outcome, which may be mediated by other measured or unmeasured factors.

As biobanks continue to grow in size and scope, new methods that are able to leverage their power while overcoming common pitfalls are required. These datasets offer unprecedented opportunity to study the causal relationship between biomarkers, complex traits and diseases. Broad analysis of the shared genetic effects of pairs of traits can be used to generate causal hypotheses that are much more likely to reflect biologically or medically-relevant phenomena than correlative analyses. It is important to point out that MR analyses without a mechanistic understanding of the biological action of each instrument are inherently speculative, with some researchers suggesting that these instead be called “joint association studies” [56]. This is especially true in large-scale analyses of noisy data, such as population-level biobanks. Indeed, we produce results that are temporally impossible; someone’s cholesterol level cannot literally cause their father’s heart disease.

Nevertheless, the interpretation of this result is clear; many of the risk alleles for cholesterol will be inherited from the father, who will have had more time to develop heart disease, resulting in high power to detect an effect. While a mechanistic understanding of the effects of each genetic instrument is ideal, there is substantial interest in the community in both applying and developing methods for causal inference using statistically associated genetic instruments. We have shown our approach is broadly useful for exploratory analysis of putatively causal effects in ever-growing databases of genotypes and phenotypes.

## 4 Methods

### Bi-direction Mendelian randomization model

We introduce a flexible model for bi-directional Mendelian randomization that encapsulates previous models for MR as special cases. In particular our model allows for both the pair of observed traits and the unobserved trait to have an arbitrary genetic architecture. It also allows for arbitrary effects of the unobserved trait on the observed traits and arbitrary effects of the observed traits on each other, including bi-directional effects. For ease of interpretation all effect sizes are modeled on a per-variance scale.

Our simulation framework (Figure 1) has 19 free parameters and 3 parameters that are a deterministic function of the others, see Table 8 for an overview. We let *N_A_* and *N_B_* represent the sample sizes of the studies for phenotypes *A* and *B*, respectively, and *M* be the number of SNPs. The effects of each phenotype on each other are given by *ν*, *η*, *γ*, and *δ* for the effect of *U* on *A*, *U* on *B*, *A* on *B* and *B* on *A*, respectively. *q*, *r*, and *s* control the proportion of variants effecting phenotypes *U*, *A*, and *B*, respectively, and 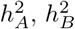 and 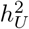. As we describe below, these parameters allow us to determine the variance of the effect size distribution for each phenotype, which we represent by 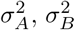 and 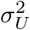. We sample the *M*-vector of effect sizes 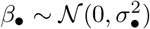 and determine which traits each SNP effects by sampling *M*-vector indicator variables *Z_•_* ∼ Bern(*π_•_*), with • representing one of *A, B* or *U*, and *π_•_* being the proportion of SNPs with non-zero direct effects on •.

We derive the variance of the phenotypes in this model. The phenotype values are given by

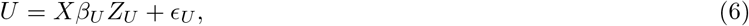

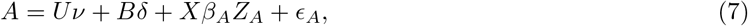

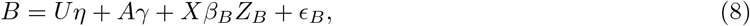

where 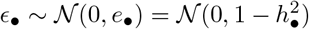 is the residual (environmental) contribution to each phenotype. Plugging the expressions for *U* and *B* into *A* and solving for *A* we find that

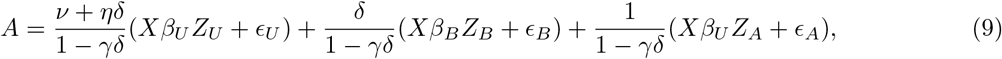

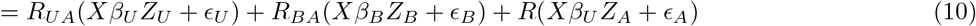

where we have introduced the shorthand *R_UA_* = (*ν* + *ηδ*)*/*(1 − *γδ*), *R_BA_* = *δ/*(1 − *γδ*) and *R* = *R_AA_* = *R_BB_* = 1*/*(1 − *γδ*) to represent the *total* causal effect of *U* on *A*, *B* on *A* and *A* on itself, respectively. It is important to notice that, because *A* can effect itself via a bi-directional effect on *B* that propagates back to *A*, *R* = 1 if and only if at least one of *γ* or *δ* is 0.

Let *g_•_* = Var(*Xβ_•_Z_•_*) be the variance component of • contributed by direct genetic effects. By mirroring the above derivation for *B*, the variance of phenotypes *A* and *B* can be broken down as

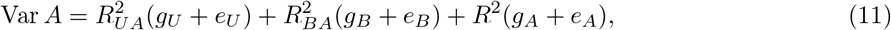

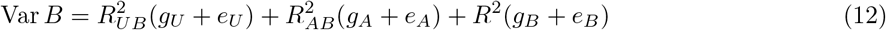

Thus if the parameters are set such that Var *A* = Var *B* = 1, then the heritabilities are given by

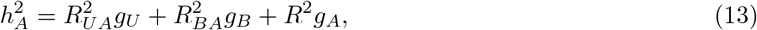

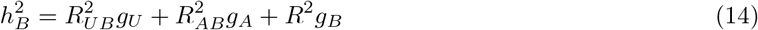

This is quite natural; the heritability of *A* is interpretable as the variance in *A* explained by *U* times the genetic component of *U*, plus the variance in *A* explained by *B* times the genetic component of *B*, plus the effect of *A* on itself times the genetic component of *A* (and likewise for *B*). The reverse is also true: if 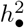 is given as above and 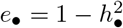, then the variances of each phenotype (Var •) are 1.

Of course, the variances of *A* and *B* will not be 1 for arbitrary settings of the parameters. Next, we show how to constrain them to be 1 so that parameters can be easily set on a per-variance scale. We manipulate (13) and (14) to determine *g_A_* and *g_B_* and thus *σ_A_* and *σ_B_*. Specifically, let

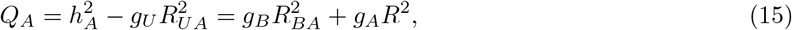

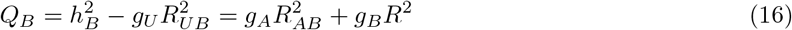

solving for *g_A_* and *g_B_* gives

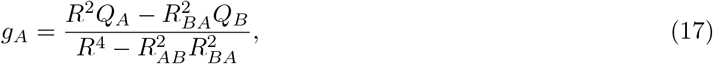

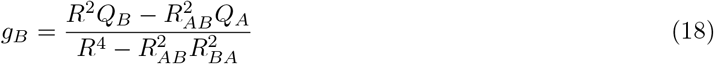

so that 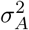 and 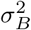 can be determined

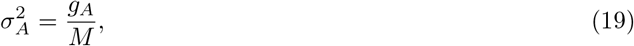

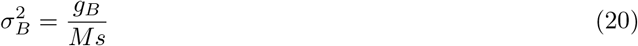

Under the null, *γ* = *δ* = 0, and therefore *R_UA_* = *ν* and *R_UB_* = *η*, while *R_BA_* = *R_AB_* = 0 and *R* = 1. In this case, the genetic covariance between *A* and *B* is 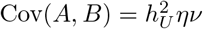. Therefore the genetic correlation is

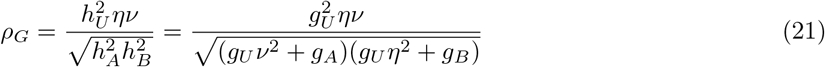

### Relationship to other models

Our model is very flexible and therefore contains other recently proposed models as special cases. Here we describe the relationship between our model and those used in LCV and CAUSE. Neither of these explicitly model the genetic architecture of the unobserved trait, preferring to tie it into the architecture of the observed traits. LCV is agnostic as to which trait is the exposure and which trait is the outcome, whereas CAUSE explicitly models one trait as the exposure (*M*, in their notation) and the other as the outcome (*Y* in their notation). For clarity when comparing to CAUSE we will use *A* as the exposure and *B* as the outcome, but it is important to remember our model is also agnostic to which trait is the exposure.

In the LCV model, under the null, the genetic correlation is *ρ_G_* = *ην*, which we can arrive at by setting 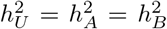. Their method attempts to quantify the deviation from a symmetric effect of the latent variable on the two observed variables, therefore the null case corresponds to *ν* = *η* (*q*_1_ = *q*_2_ in their notation). Finally, their settings focus on the case where the effect distribution of the SNPs acting on the observed traits is the same for SNPs acting directly and via the latent variable. In our model we can enact this assumption via the constraints 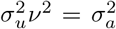 and 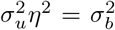. Finally, we assume that the SNPs effect a single trait in expectation, that is *q* + *r* + *s* = 1. Under the null, 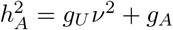. Using the assumption that 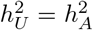, we have that 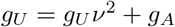. Rearranging and simplifying, we have 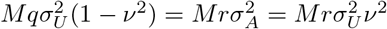 and thus *r* = *q*(1 − *ν*2)*/ν*2 (and likewise for *s*). Plugging into *q* + *r* + *s* = 1 and simplifying leads to

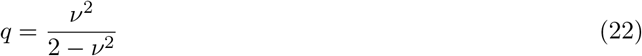

The CAUSE model also assumes that 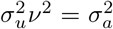. This is represented by the 1 on the arrow from the *u a* latent factor to the exposure in their model, indicating that SNPs have the same effect distribution on the exposure when acting via the latent variable or directly. In that model, the proportion of variants effecting the unobserved variable *q* controls the magnitude of the genetic correlation which then implicitly determines the heritibility of the hidden variable 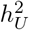. Again using 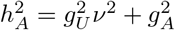 and substituting 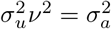 we find 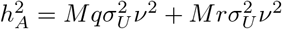. Solving for 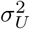 and using the fact that 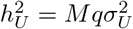 we find

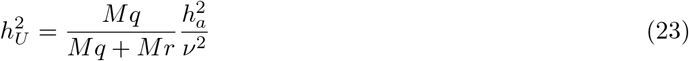

## 5 Data Availability

UK Biobank summary statistics are available from the Neal lab at https://www.nealelab.is/uk-biobank. Our full data analysis results are available at https://zenodo.org/record/4605239.

## 6 Code Availability

All code used in the production of this manuscript is available at https://github.com/brielin/WWER.

## 7 Acknowledgments

BCB and DAK would like to thank Tuuli Lappalainen for helpful feedback on the manuscript. BCB is funded by post-doctoral fellowship from the Data Science Institute at Columbia University.

## 8 Author Contributions

BCB and DAK jointly formulated the model and estimation procedure. BCB wrote the code, conducted analyses, and drafted the manuscript. DAK supervised and assisted with editing the manuscript.

---

**Table S1:** A list of all parameters in our model, with examples and feasible ranges.

| Symbol | Definition | Class | Possible values | Example |
| --- | --- | --- | --- | --- |
| $M$ | Number of SNPs | Fixed | $[1, \infty)$ | 500,000 |
| $N_A$ | Sample size of study for phenotype A | Fixed | $[1, \infty)$ | 50,000 |
| $N_B$ | Sample size of study for phenotype B | Fixed | $[1, \infty)$ | 50,000 |
| $\nu$ | Per-variance effect of U on A | Fixed | $[-1, 1]$ | $\sqrt{0.2}$ |
| $\eta$ | Per-variance effect of U on B | Fixed | $[-1, 1]$ | $\sqrt{0.2}$ |
| $\gamma$ | Per-variance effect of A on B | Fixed | $[-1, 1]$ | $\sqrt{0.2}$ |
| $\delta$ | Per-variance effect of B on A | Fixed | $[-1, 1]$ | $\sqrt{0.2}$ |
| $q$ | Proportion of variants effecting U | Fixed | $[0, 1]$ | 0.3 |
| $r$ | Proportion of variants effecting A | Fixed | $[0, 1]$ | 0.3 |
| $s$ | Proportion of variants effecting B | Fixed | $[0, 1]$ | 0.3 |
| $h_A^2$ | Heritability of phenotype A | Fixed | $[0, 1]$ | 0.2 |
| $h_B^2$ | Heritability of phenotype B | Fixed | $[0, 1]$ | 0.2 |
| $h_U^2$ | Heritability of phenotype U | Fixed | $[0, 1]$ | 0.2 |
| $\sigma_A$ | Variance of the effect size distribution for A | Fixed | NA | $1e-4$ |
| $\sigma_B$ | Variance of the effect size distribution for B | Fixed | NA | $1e-4$ |
| $\sigma_U$ | Variance of the effect size distribution for U | Fixed | NA | $1e-4$ |
| $\beta_A$ | Effect of SNP on phenotype A, $\sim \mathcal{N}(0, \sigma_A)$ | Random | $[-1, 1]$ | $\sqrt{0.01}$ |
| $\beta_B$ | Effect of SNP on phenotype B, $\sim \mathcal{N}(0, \sigma_A)$ | Random | $[-1, 1]$ | $\sqrt{0.01}$ |
| $\beta_U$ | Effect of SNP on phenotype U, $\sim \mathcal{N}(0, \sigma_A)$ | Random | $[-1, 1]$ | $\sqrt{0.01}$ |
| $Z_A$ | Indicator that SNP effects A, sampled from $Bern(r)$ | Random | $\{0, 1\}$ | 0 |
| $Z_B$ | Indicator that SNP effects B, sampled from $Bern(s)$ | Random | $\{0, 1\}$ | 0 |
| $Z_U$ | Indicator that SNP effects U, sampled from $Bern(q)$ | Random | $\{0, 1\}$ | 0 |

**Figure S1:**
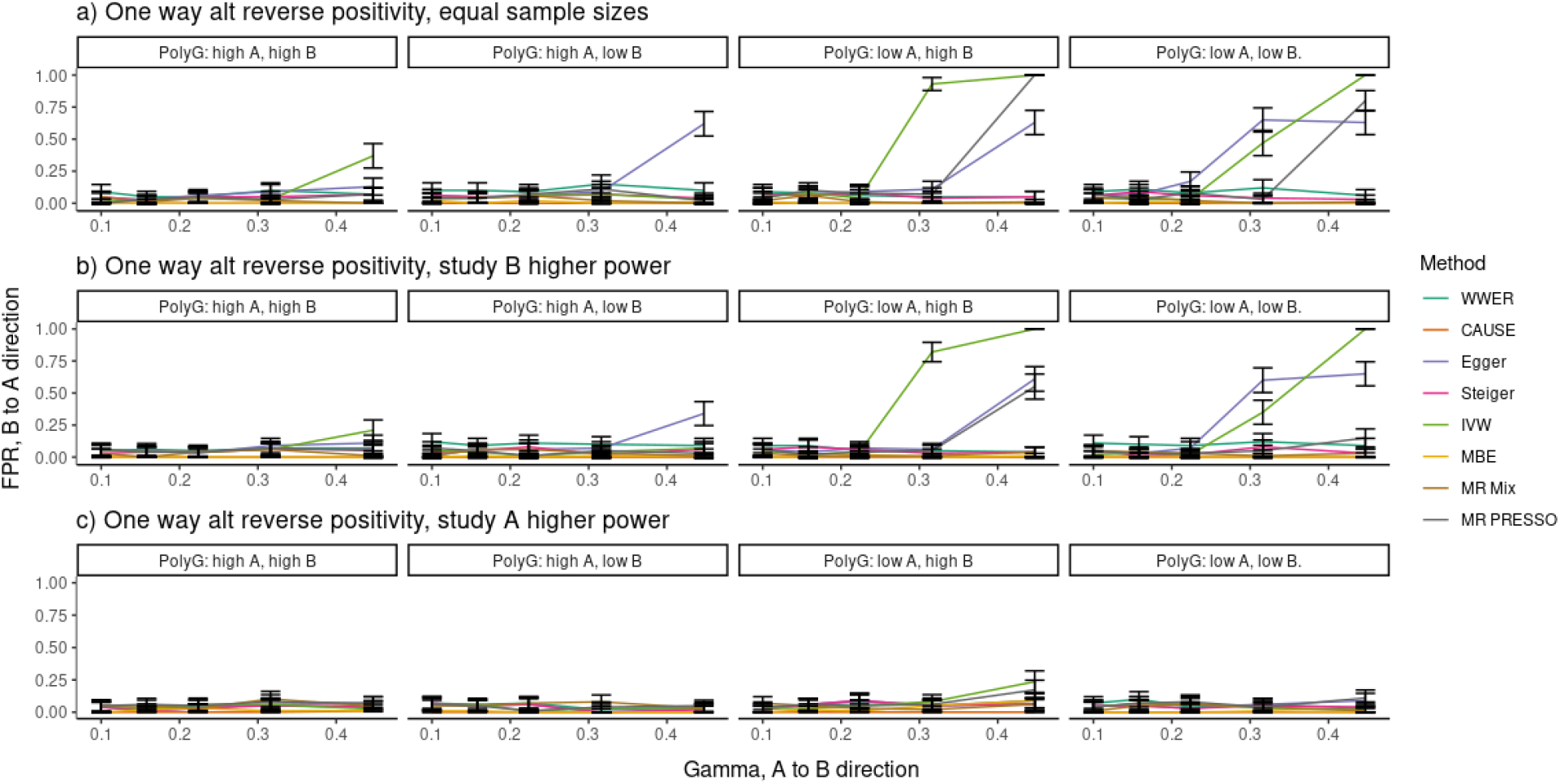
False positive rate in the reverse direction (B to A) when there is a causal effect in the A to B direction (x-axis). Settings correspond to the settings of Figure 3a-c. IVW, MR Presso and Egger often produce a false positive in the B to A direction when there is a strong effect in the A to B direction.

**Figure S2:**
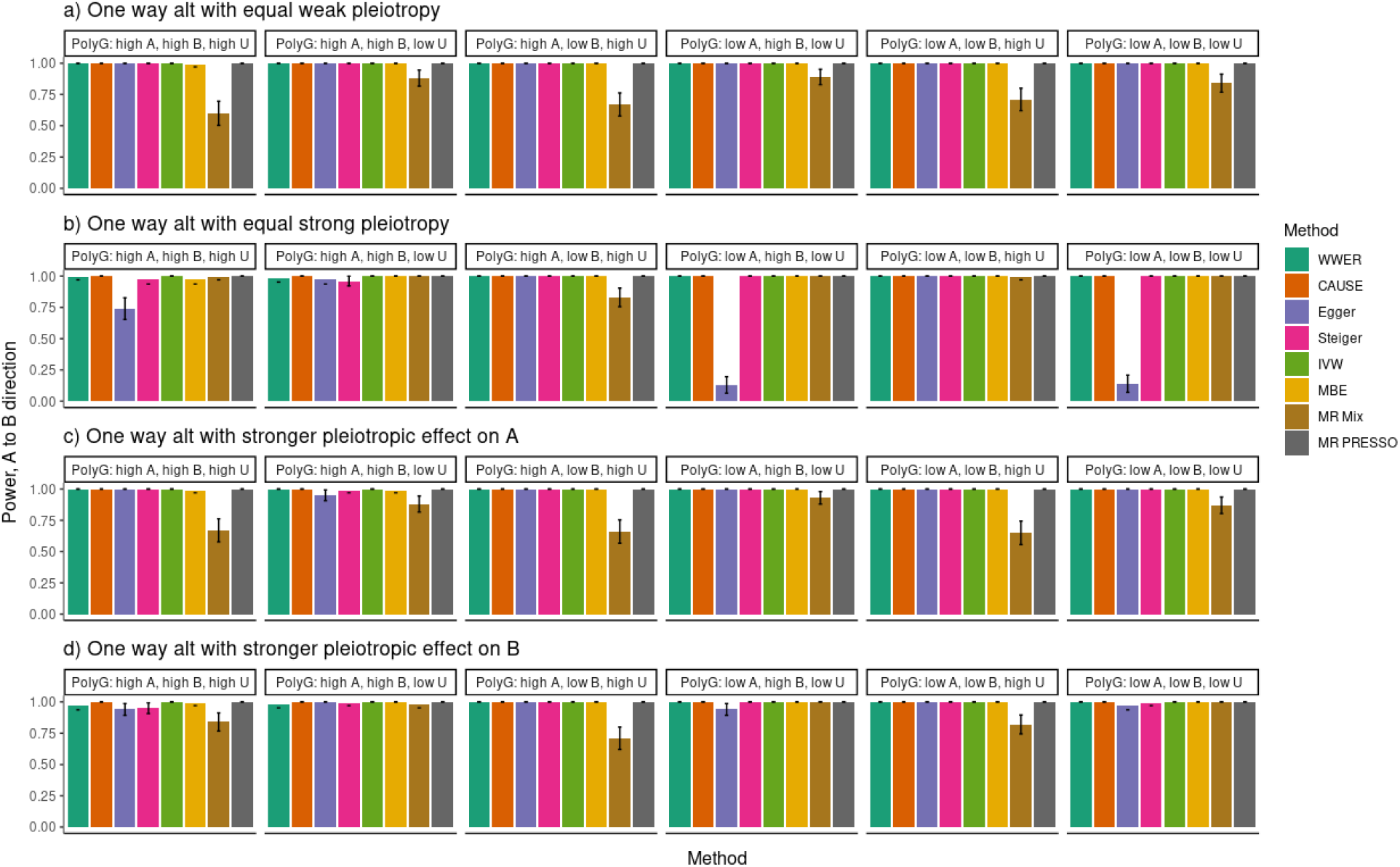
Power of each method for various settings when there is both a causal effect and correlated pleiotropy.

**Figure S3:**
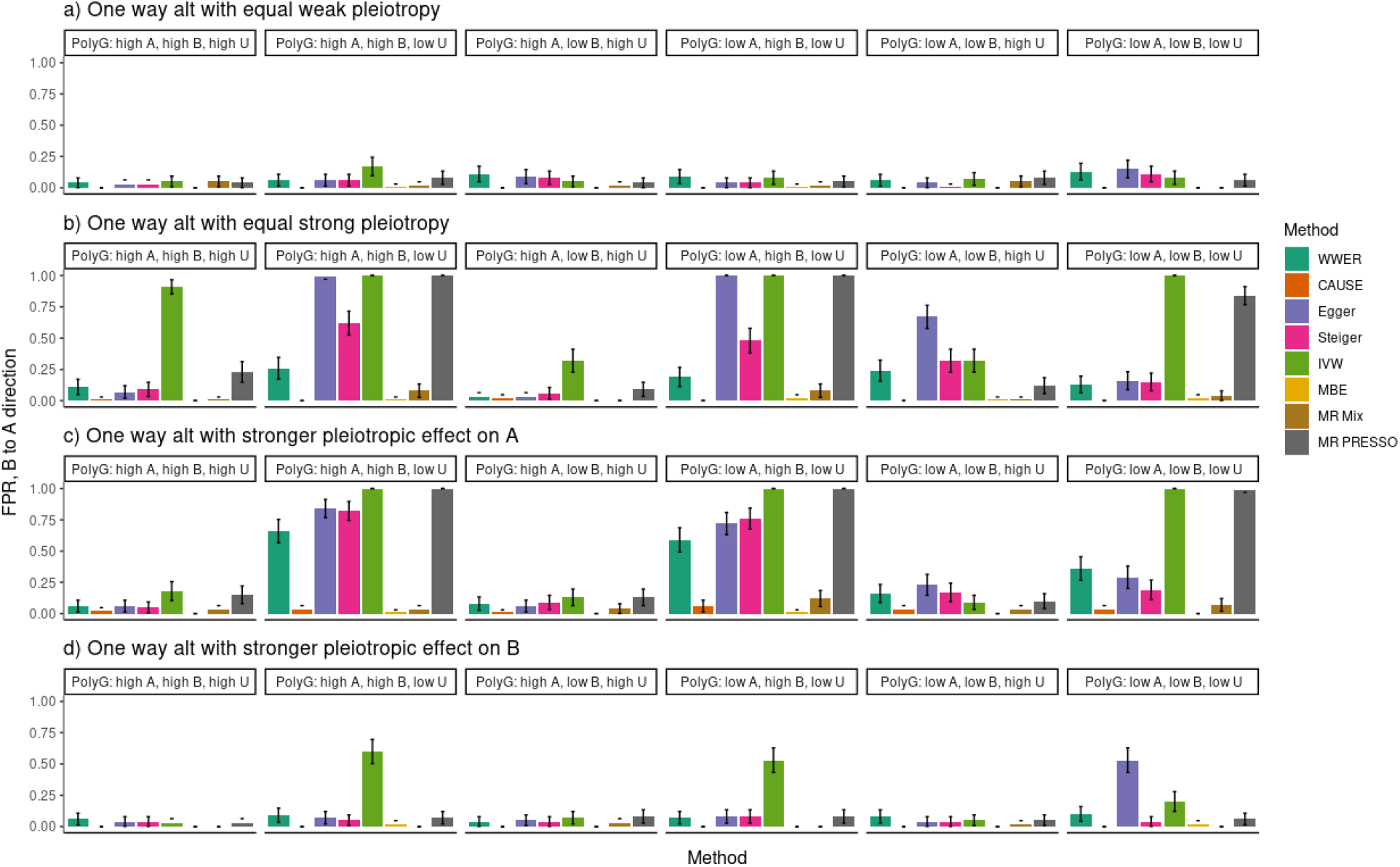
False positive rate in the reverse direction (B to A) when there is a causal effect in the forward direction (A to B) and correlated pleiotropy. Settings correspond to the settings in Figure S2. Both WWER and Stieger filtering show a substantial improvement over Egger regression and other traditional methods. Here WWER clearly outperforms Steiger filtering in several settings, although there are still two where it produces a high FPR.

**Figure S4:**
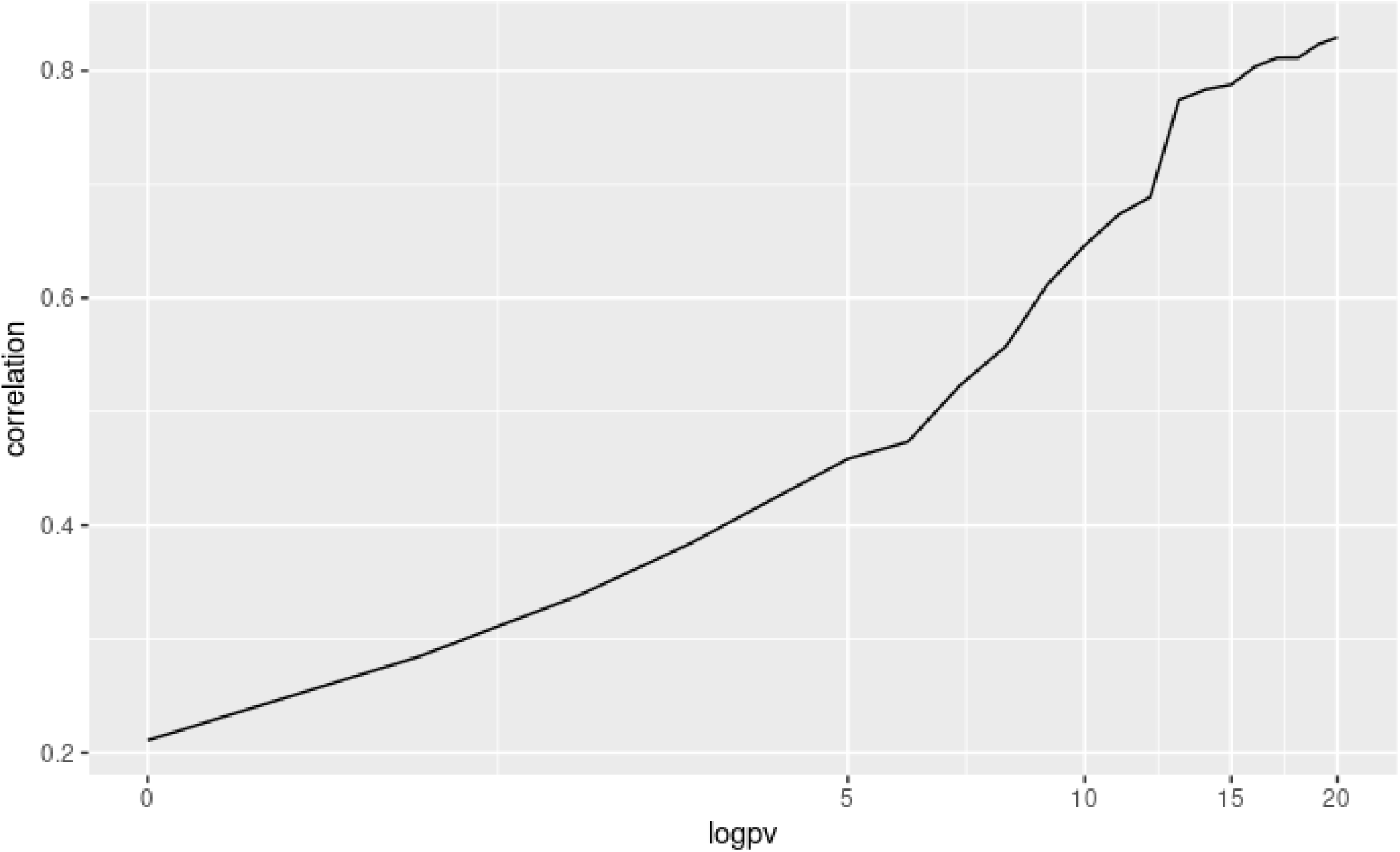
Correlation of the estimated causal effect with the genetic correlation, as a function of the significance of the estimated causal effect. The global correlation is low, but when the estimate of the causal effect is more significant it is also more similar to the estimated genetic correlation.

## References

[1] Jack Bowden, George Davey Smith, and Stephen Burgess. “Mendelian randomization with invalid instruments: Effect estimation and bias detection through Egger regression”. In: International Journal of Epidemiology 44.2 (2015), pp. 512–525. issn: 14643685. doi: 10.1093/ije/dyv080.

[2] Fernando Pires Hartwig, George Davey Smith, and Jack Bowden. “Robust inference in summary data Mendelian randomization via the zero modal pleiotropy assumption”. In: International Journal of Epidemiology 46.6 (2017), pp. 1985–1998. issn: 14643685. doi: 10.1093/ije/dyx102.

[3] Jean Morrison et al. “Mendelian randomization accounting for correlated and uncorrelated pleiotropic effects using genome-wide summary statistics”. In: Nature Genetics 52.7 (July 2020), pp. 740–747. issn: 15461718. doi: 10.1038/s41588-020-0631-4. url: https://doi.org/10.1038/s41588-020-0631-4.

[4] Guanghao Qi and Nilanjan Chatterjee. “Mendelian randomization analysis using mixture models for robust and efficient estimation of causal effects”. In: Nature Communications 10.1 (Dec. 2019), pp. 1–10. issn: 20411723. doi: 10.1038/s41467-019-09432-2. url: https://doi.org/10.1038/s41467-019-09432-2.

[5] Luke J O’Connor and Alkes L Price. “Distinguishing genetic correlation from causation across 52 diseases and complex traits”. In: Nature Genetics 50.12 (2018), pp. 1728–1734. issn: 1546-1718. doi: 10.1038/s41588-018-0255-0. url: https://doi.org/10.1038/s41588-018-0255-0.

[6] N. J. Timpson et al. “C-reactive protein levels and body mass index: Elucidating direction of causation through reciprocal Mendelian randomization”. In: International Journal of Obesity 35.2 (2011), pp. 300–308. issn: 03070565. doi: 10.1038/ijo.2010.137.

[7] Rebecca C. Richmond et al. “Assessing Causality in the Association between Child Adiposity and Physical Activity Levels: A Mendelian Randomization Analysis”. In: PLoS Medicine 11.3 (2014). issn: 15491676. doi: 10.1371/journal.pmed.1001618. url: https://pubmed.ncbi.nlm.nih.gov/24642734/.

[8] Joseph K. Pickrell et al. “Detection and interpretation of shared genetic influences on 42 human traits”. In: Nature Genetics 48.7 (2016), pp. 709–717. issn: 15461718. doi: 10.1038/ng.3570. url: http://dx.doi.org/10.1038/ng.3570.

[9] Gibran Hemani et al. “Automating Mendelian randomization through machine learning to construct a putative causal map of the human phenome”. In: (2017). doi: 10.1101/173682. url: https://doi.org/10.1101/173682.

[10] B. L. welch. “The generalisation of student’s problems when several different population variances are involved.” In: Biometrika 34.1-2 (1947), pp. 28–35. issn: 00063444. doi: 10.1093/biomet/34.1-2.28. url: https://pubmed.ncbi.nlm.nih.gov/20287819/.

[11] Marie Verbanck et al. “Detection of widespread horizontal pleiotropy in causal relationships inferred from Mendelian randomization between complex traits and diseases”. In: Nature Genetics 50.5 (May 2018), pp. 693–698. issn: 15461718. doi: 10.1038/s41588-018-0099-7. url: https://doi.org/10.1038/s41588-018-0099-7.

[12] Qingyuan Zhao et al. “Statistical inference in two-sample summary-data Mendelian randomization using robust adjusted profile score”. In: Annals of Statistics 48.3 (June 2020), pp. 1742–1769. issn: 21688966. doi: 10.1214/19-AOS1866. arXiv: 1801.09652. url: https://projecteuclid.org/euclid.aos/1594972837.

[13] Jack Bowden et al. “Consistent Estimation in Mendelian Randomization with Some Invalid Instruments Using a Weighted Median Estimator”. In: Genetic Epidemiology 40.4 (May 2016), pp. 304–314. issn: 10982272. doi: 10.1002/gepi.21965. url: https://pubmed.ncbi.nlm.nih.gov/27061298/.

[14] Michael R. Shurin and Yuri S. Smolkin. “Immune-mediated diseases: Where do we stand?” In: Advances in Experimental Medicine and Biology. Vol. 601. 2007, pp. 3–12. isbn: 9780387720043. doi: 10.1007/978-0-387-72005-0_1.

[15] Nasa Sinnott-Armstrong et al. “Genetics of 35 blood and urine biomarkers in the UK Biobank”. In: Nature Genetics (2021). 1061-4036. doi: 10.1038/s41588-020-00757-z. url: http://dx.doi.org/10.1038/s41588-020-00757-z.

[16] Clare Bycroft et al. “The UK Biobank resource with deep phenotyping and genomic data”. In: Nature 562.7726 (Oct. 2018), pp. 203–209. 14764687. doi: 10.1038/s41586-018-0579-z. url: https://doi.org/10.1038/s41586-018-0579-z.

[17] UK Biobank — Neale lab. url: http://www.nealelab.is/uk-biobank (visited on 04/09/2021).

[18] M Ellaurie. “Platelet abnormalities in asthma and allergy*1”. In: Journal of Allergy and Clinical Immunology 113.2 (Feb. 2004), S161. 00916749. doi: 10.1016/j.jaci.2004.01.009.

[19] Paul Stoll and Marek Lommatzsch. “Platelets in Asthma: Does Size Matter?” In: Respiration 88.1 (2014), pp. 22–23. 0025-7931. doi: 10.1159/000362798. url: https://www.karger.com/Article/FullText/362798.

[20] ManalR Hafez et al. “Assessment of bronchial asthma exacerbation: the utility of platelet indices”. In: Egyptian Journal of Bronchology 13.5 (Jan. 2019), p. 623. 1687-8426. doi: 10.4103/ejb.ejb_69_19. url: http://www.ejbronchology.eg.net/text.asp?2019/13/5/623/276301.

[21] John W. Semple, Joseph E. Italiano, and John Freedman. Platelets and the immune continuum. Apr. 2011. doi: 10.1038/nri2956. url: https://pubmed.ncbi.nlm.nih.gov/21436837/.

[22] Fauzia Imtiaz et al. “Neutrophil lymphocyte ratio as a measure of systemic inflammation in prevalent chronic diseases in Asian population”. In: International Archives of Medicine 5.1 (2012). 17557682. doi: 10.1186/1755-7682-5-2. url: https://pubmed.ncbi.nlm.nih.gov/22281066/.

[23] Özlem Taşoğlu et al. “Is blood neutrophil-lymphocyte ratio an independent predictor of knee os-teoarthritis severity?” In: Clinical Rheumatology 35.6 (June 2016), pp. 1579–1583. 14349949. doi: 10.1007/s10067-016-3170-8. url: https://pubmed.ncbi.nlm.nih.gov/26780447/.

[24] Adil Can Gungen and Yusuf Aydemir. The correlation between asthma disease and neutrophil to lymphocyte ratio. Tech. rep. 0. 2017. url: http://www.alliedacademies.org/research-journal-of-allergy-and-immunology/.

[25] Sophie Demarche et al. “Detailed analysis of sputum and systemic inflammation in asthma phenotypes: Are paucigranulocytic asthmatics really non-inflammatory?” In: BMC Pulmonary Medicine 16.1 (Apr. 2016), pp. 1–13. 14712466. doi: 10.1186/s12890-016-0208-2. url: https://link.springer.com/articles/10.1186/s12890-016-0208-2%20https://link.springer.com/article/10.1186/s12890-016-0208-2.

[26] Ruurt A. Jukema, Tarek A.N. Ahmed, and Jean Claude Tardif. “Does low-density lipoprotein cholesterol induce inflammation? if so, does it matter? Current insights and future perspectives for novel therapies”. In: BMC Medicine 17.1 (Nov. 2019), p. 197. 17417015. doi: 10.1186/s12916-019-1433-3. url:https://bmcmedicine.biomedcentral.com/articles/10.1186/s12916-019-1433-3.

[27] Alexandros Tsoupras, Ronan Lordan, and Ioannis Zabetakis. Inflammation, not cholesterol, is a cause of chronic disease. May 2018. doi: 10.3390/nu10050604. url: pmc/articles/PMC5986484/%20/pmc/articles/PMC5986484/?report=abstract%20https://www.ncbi.nlm.nih.gov/pmc/articles/PMC5986484/.

[28] M. B. Fessler et al. “Relationship of serum cholesterol levels to atopy in the US population”. In: Allergy: European Journal of Allergy and Clinical Immunology 65.7 (2010), pp. 859–864. 13989995. doi: 10.1111/j.1398-9995.2009.02287.x. url: /pmc/articles/PMC4045486/%20/pmc/articles/PMC4045486/?report=abstract%20https://www.ncbi.nlm.nih.gov/pmc/articles/PMC4045486/.

[29] William J. Calhoun, JULIE Sedgwick, and William W. Busse. “The Role of Eosinophils in the Pathophysiology of Asthma”. In: Annals of the New York Academy of Sciences 629.1 Advances in t (July 1991), pp. 62–72. 0077-8923. doi: 10.1111/j.1749-6632.1991.tb37961.x. url: http://doi.wiley.com/10.1111/j.1749-6632.1991.tb37961.x.

[30] Dirk Smith et al. “A rare IL33 loss-of-function mutation reduces blood eosinophil counts and protects from asthma”. In: PLOS Genetics 13.3 (Mar. 2017). Ed. by Tuuli Lappalainen, e1006659. 1553-7404. doi: 10.1371/journal.pgen.1006659. url: https://dx.plos.org/10.1371/journal.pgen.1006659.

[31] Maria Cappello et al. Liver function test abnormalities in patients with inflammatory bowel diseases: A hospital-based survey. June 2014. doi: 10.4137/CGast.S13125. url: /pmc/articles/PMC4069044/%20/pmc/articles/PMC4069044/?report=abstract%20https://www.ncbi.nlm.nih.gov/pmc/articles/PMC4069044/.

[32] Jessika Barendregt et al. “Liver test abnormalities predict complicated disease behaviour in patients with newly diagnosed Crohn’s disease”. In: International Journal of Colorectal Disease 32.4 (Apr. 2017), pp. 459–467. 14321262. doi: 10.1007/s00384-016-2706-3. url: https://link.springer.com/article/10.1007/s00384-016-2706-3.

[33] Emma H. Baker and Derek Bell. Blood glucose: Of emerging importance in COPD exacerbations. Oct. 2009. doi: 10.1136/thx.2009.118638. url: http://thorax.bmj.com/.

[34] Michael J. Berridge. Vitamin D deficiency and diabetes. Apr. 2017. doi: 10.1042/BCJ20170042. url: https://pubmed.ncbi.nlm.nih.gov/28341729/.

[35] P. Rocha-Pereira et al. “Erythrocyte damage in mild and severe psoriasis”. In: British Journal of Dermatology 150.2 (Feb. 2004), pp. 232–244. 00070963. doi: 10.1111/j.1365-2133.2004.05801. x. url: https://pubmed.ncbi.nlm.nih.gov/14996093/.

[36] Susana Coimbra et al. “The roles of cells and cytokines in the pathogenesis of psoriasis”. In: International Journal of Dermatology 51.4 (Apr. 2012), pp. 389–398. 00119059. doi: 10.1111/j.1365-4632.2011.05154.x. url: http://doi.wiley.com/10.1111/j.1365-4632.2011.05154.x.

[37] Sibel Doğan and Nilgün Atakan. “Red blood cell distribution width is a reliable marker of inflammation in plaque psoriasis”. In: Acta Dermatovenerologica Croatica 25.1 (2017), pp. 26–31. 18476538.

[38] C. Vahlquist, G. Michaelsson, and B. Vessby. “Serum lipoproteins in middle-aged men with psoriasis”. In: Acta Dermato-Venereologica 67.1 (1987), pp. 12–15. 00015555.

[39] Mehdi Taheri Sarvtin et al. “Study of serum lipids and lipoproteins in patients with psoriasis”. In: Journal of Mazandaran University of Medical Sciences 23.98 (2013), pp. 173–177. 17359260.

[40] Michael Holzer et al. “Psoriasis alters HDL composition and cholesterol efflux capacity”. In: Journal of Lipid Research 53.8 (Aug. 2012), pp. 1618–1624. 00222275. doi: 10.1194/jlr.M027367. url: http://www.jlr.org.

[41] V. Sathiyapriya et al. “Evidence for the role of lipid peroxides on glycation of hemoglobin and plasma proteins in non-diabetic asthma patients”. In: Clinica Chimica Acta 366.1-2 (Apr. 2006), pp. 299–303. 00098981. doi: 10.1016/j.cca.2005.11.001.

[42] Miyoshi Fujita et al. “C-reactive protein levels in the serum of asthmatic patients”. In: Annals of Allergy, Asthma and Immunology 99.1 (July 2007), pp. 48–53. 10811206. doi: 10.1016/S1081-1206(10)60620-5.

[43] Natalie M. Niessen et al. “Neutrophilic asthma features increased airway classical monocytes”. In: Clinical & Experimental Allergy 51.2 (Feb. 2021), pp. 305–317. 0954-7894. doi: 10.1111/cea.13811. url: https://onlinelibrary.wiley.com/doi/10.1111/cea.13811.

[44] Ibon Eguìluz-Gracia et al. “Monocytes accumulate in the airways of children with fatal asthma”. In: Clinical and Experimental Allergy 48.12 (Dec. 2018), pp. 1631–1639. 13652222. doi: 10.1111/cea.13265. url: https://pubmed.ncbi.nlm.nih.gov/30184280/.

[45] A. Barry Kay. The role of T lymphocytes in asthma. 2006. doi: 10.1159/000090230. url: https://pubmed.ncbi.nlm.nih.gov/16354949/.

[46] Telugu A. Narasaraju et al. “Expression profile of IGF system during lung injury and recovery in rats exposed to hyperoxia: A possible role of IGF-1 in alveolar epithelial cell proliferation and differentiation”. In: Journal of Cellular Biochemistry 97.5 (Apr. 2006), pp. 984–998. 0730-2312. doi: 10.1002/jcb.20653. url: http://doi.wiley.com/10.1002/jcb.20653.

[47] Yueh Ying Han et al. “Serum insulin-like growth factor-1, asthma, and lung function among British adults”. In: Annals of Allergy, Asthma and Immunology 126.3 (Mar. 2021), 284–291.e2. 15344436. doi: 10.1016/j.anai.2020.12.005.

[48] Mojtaba Shafiee et al. “Depression and anxiety symptoms are associated with white blood cell count and red cell distribution width: A sex-stratified analysis in a population-based study”. In: Psychoneuroendocrinology 84 (Oct. 2017), pp. 101–108. 18733360. doi: 10.1016/j.psyneuen.2017.06.021. url: https://pubmed.ncbi.nlm.nih.gov/28697416/.

[49] Yuji Shimizu et al. “Short stature is an inflammatory disadvantage among middle-aged Japanese men”. In: Environmental Health and Preventive Medicine 21.5 (Sept. 2016), pp. 361–367. 13474715. doi: 10.1007/s12199-016-0538-y. url: /pmc/articles/PMC5305990/%20/pmc/articles/PMC5305990/?report=abstract%20https://www.ncbi.nlm.nih.gov/pmc/articles/PMC5305990/.

[50] P. S. Sanmuganathan et al. “Aspirin for primary prevention of coronary heart disease: Safety and absolute benefit related to coronary risk derived from meta-analysis of randomised trials”. In: Heart 85.3 (Mar. 2001), pp. 265–271. 13556037. doi: 10.1136/heart.85.3.265. url: www.heartjnl.com.

[51] Thomas M. MacDonald and Li Wei. “Is there an Interaction between the Cardiovascular Protective Effects of Low-Dose Aspirin and Ibuprofen?” In: Basic ¡html ent glyph=”@amp;” ascii=”&”/¿ Clinical Pharmacology ¡html ent glyph=”@amp;” ascii=”&”/¿ Toxicology 98.3 (Mar. 2006), pp. 275–280. 1742-7835. doi: 10.1111/j.1742-7843.2006.pto_371.x. url: http://doi.wiley.com/10.1111/j.1742-7843.2006.pto%7B%5C_%7D371.x.

[52] Yanyan Zhu, Yuqing Zhang, and Hyon K. Choi. “The serum urate-lowering impact of weight loss among men with a high cardiovascular risk profile: The Multiple Risk Factor Intervention Trial”. In: Rheumatology 49.12 (Dec. 2010), pp. 2391–2399. 14620324. doi: 10.1093/rheumatology/keq256. url: https://pubmed.ncbi.nlm.nih.gov/20805117/.

[53] Marjolein Visser et al. “Elevated C-reactive protein levels in overweight and obese adults”. In: Journal of the American Medical Association 282.22 (Dec. 1999), pp. 2131–2135. 00987484. doi: 10.1001/jama.282.22.2131. url: https://pubmed.ncbi.nlm.nih.gov/10591334/.

[54] Rana H. Mosli and Hala H. Mosli. “Obesity and morbid obesity associated with higher odds of hypoalbuminemia in adults without liver disease or renal failure”. In: Diabetes, Metabolic Syndrome and Obesity: Targets and Therapy 10 (Nov. 2017), pp. 467–472. 11787007. doi: 10.2147/DMSO.S149832. url: /pmc/articles/PMC5687480/%20/pmc/articles/PMC5687480/?report=abstract%20https://www.ncbi.nlm.nih.gov/pmc/articles/PMC5687480/.

[55] Bogdan Pasaniuc and Alkes L. Price. Dissecting the genetics of complex traits using summary association statistics. Feb. 2017. doi: 10.1038/nrg.2016.142. url: /pmc/articles/PMC5449190/%20/pmc/articles/PMC5449190/?report=abstract%20https://www.ncbi.nlm.nih.gov/pmc/articles/PMC5449190/.

[56] Stephen Burgess, Adam S. Butterworth, and John R. Thompson. “Beyond Mendelian randomization: How to interpret evidence of shared genetic predictors”. In: Journal of Clinical Epidemiology 69 (Jan. 2016), pp. 208–216. 18785921. doi: 10.1016/j.jclinepi.2015.08.001. url: /pmc/articles/PMC4687951/%20/pmc/articles/PMC4687951/?report=abstract%20https://www.ncbi.nlm.nih. gov/pmc/articles/PMC4687951/.

